# Gut metagenomes of Asian octogenarians reveal metabolic potential expansion and distinct microbial species associated with aging phenotypes

**DOI:** 10.1101/2024.07.08.602612

**Authors:** Aarthi Ravikrishnan, Indrik Wijaya, Eileen Png, Kern Rei Chng, Eliza Ho Xin Pei, Amanda Ng Hui Qi, Ahmad Nazri Mohamed Naim, Jean-Sebastien Gounot, Guan Shou Ping, Hanqing Jasinda Lee, Guan Lihuan, Li Chenhao, Jayce Koh Jia Yu, Paola Florez de Sessions, Woon-Puay Koh, Lei Feng, Tze Pin Ng, Anis Larbi, Andrea B. Maier, Brian Kennedy, Niranjan Nagarajan

**Author notes:** Joint First Authors (Aarthi Ravikrishnan, Indrik Wijaya).

## Abstract

While rapid demographic changes in Asia are driving the incidence of chronic diseases related to aging, the limited availability of high-quality *in vivo* data hampers our ability to understand complex multi-factorial contributions, including gut microbial, to healthy aging. Leveraging the availability of a well-phenotyped cohort of community-living octogenarians in Singapore, we used deep shotgun metagenomic sequencing to do high-resolution taxonomic and functional characterization of their gut microbiomes (n=234). Joint species-level analysis with other Asian cohorts identified a distinct age-associated shift in Asian gut metagenomes, characterized by a reduction in microbial richness, and enrichment of specific *Alistipes* and *Bacteroides* species (e.g. *Alistipes shahii* and *Bacteroides xylanisolvens*). Functional pathway analysis confirmed that these changes correspond to a metabolic potential expansion in aging towards alternate pathways that synthesize and utilize amino-acid precursors, relative to the dominant microbial guilds that typically produce butyrate in the gut from pyruvate (e.g. *Faecalibacterium prausnitzii, Roseburia inulinivorans*). Extending these observations to key clinical markers helped identify >10 robust gut microbial associations to inflammation, cardiometabolic and liver health, including potential probiotic species such as *Parabacteroides goldsteinii* and pathobionts such as *Klebsiella pneumoniae*, highlighting the role of the microbiome as biomarkers and potential intervention targets for promoting healthy aging.

## Introduction

Over the last few decades, economic changes and advances in healthcare systems have significantly improved life expectancies in Asia^1, 2^, and are expected to lead to a rapid shift in demographics by doubling the population of 60+ individuals by 2050^3^ (from 19.7% in 2015 to 40%). This has been accompanied by a rising incidence of chronic diseases linked to aging^4, 5^, and corresponding socio-economic stress on healthcare systems across Asia^6, 7^. There is therefore an urgent need to identify lifestyle, dietary and pharmaceutical interventions that promote healthy aging in Asian populations^8–10^.

Aging is believed to be a complex, multi-factorial phenomenon with progressive decline in several physiological functions including in the gastrointestinal and immune system^11, 12^. Not surprisingly, many studies have therefore identified correlations between gut microbiome composition and age^13, 14^. For example, in recent work, Zhang et al^15^ have shown that there are distinct age and gender-associated trajectories in the gut microbiome that are consistent across populations. Several studies have also highlighted that frailty indices and healthy aging are linked to the state of the gut microbiome^16, 17^, along with known gut-microbiome associations with aging-related diseases^18, 19^. In particular, shifts in several families and genera of bacteria including *Bacteroidetes, Proteobacteria, Roseburia* and *Escherichia* have been associated via 16S rRNA sequencing to healthy aging^20–26^. Work by Wilmanski et al also suggests that compositional uniqueness of the gut microbiome may serve as a marker for healthy aging^27^. To compensate for the potential low resolution of 16S studies, recent work has sought to leverage shotgun metagenomic sequencing to characterize the gut microbiome in elderly populations, but have been limited by cohort size (n<50) and the spectrum of age ranges that they study, in their ability to account for potential confounders while identifying microbial mechanisms associated with healthy aging^28, 29^.

To address this, we used deep shotgun metagenomic analysis (n=234, >20 million reads on average) to study gut microbiomes in a cohort of community-living octogenarians (primarily, age range=[71-100], **Table 1**) in Singapore^30^. The gut microbiomes of healthy octogenarians exhibited a defined shift in diversity and overall taxonomic composition, as a function of age and independent of other covariates. In addition, taxonomic analysis identified several species-level changes (e.g. enrichment of *Alistipes shahii* and *Bacteroides xylanisolvens*), consistent with functional pathway analysis of the data, that points to a metabolic potential expansion in aging towards alternate pathways that synthesize and utilize amino-acids (e.g. L-lysine) as precursor substrates, relative to the classical pyruvate-to-butyrate production pathway. As butyrate derived from gut bacteria has diverse roles in host health (e.g. as energy for colonocytes, and reducing gut inflammation^31–33^), we next associated gut microbiome composition with key markers for inflammation (e.g. CRP), cardiometabolic health (e.g. fasting blood glucose) and liver health (e.g. AST), to find additional microbial functions associated with healthy aging. Leveraging extensive clinical data and the size of the metagenomic datasets, we identified >10 robust associations (accounting for demographic and clinical covariates that can be confounders) that highlight the role of the microbiome as biomarkers and potential intervention targets for promoting healthy aging.

**Table 1:** Clinical characteristics of the SG90 cohort.

|  | All Subjects | Male | Female |
| --- | --- | --- | --- |
| **Demographics** |  |  |  |
| Number of subjects | 234 | 95 | 139 |
| Age (Mean ± SD) | 86 ± 4 | 86 ± 3 | 86 ± 4 |
| **Serum measurements (Mean ± SD)** |  |  |  |
| High-sensitivity CRP (mg/L) | 3.7 ± 8.4 | 4.6 ± 11.9 | 3.1 ± 4.4 |
| Fasting blood glucose (mmol/L) | 5.2 ± 0.4 | 5.3 ± 0.5 | 5.2 ± 0.4 |
| Triglycerides (mmol/L) | 1.4 ± 0.6 | 1.2 ± 0.6 | 1.5 ± 0.7 |
| Total cholesterol (mmol/L) | 4.6 ± 1.1 | 4.4 ± 1.0 | 4.8 ± 1.1 |
| High-density lipoprotein (mmol/L) | 1.4 ± 0.3 | 1.3 ± 0.3 | 1.4 ± 0.4 |
| Low-density lipoprotein (mmol/L) | 2.6 ± 0.9 | 2.5 ± 0.8 | 2.7 ± 0.9 |
| Alanine aminotransferase (U/L) | 18.4 ± 8.0 | 19.3 ± 8.1 | 17.8 ± 8.0 |
| Aspartate aminotransferase (U/L) | 25.0 ± 6.8 | 25.5 ± 7.6 | 24.6 ± 6.2 |
| **Disease incidence (%)** |  |  |  |
| Type 2 diabetes | 18 | 12 | 22 |
| Heart attack | 6 | 9 | 5 |
| Heart failure | 4 | 4 | 4 |
| Cataracts/glaucoma | 36 | 31 | 38 |
| Kidney failure | 0.9 | 1 | 0.7 |
| Asthma | 3 | 1 | 4 |
| COPD | 2 | 4 | 0.7 |
| Arthritis | 9 | 5 | 12 |
| Osteoporosis | 4 | 1 | 6 |
| Hip fracture | 4 | 1 | 6 |
| Parkinson's disease | 0.9 | 2 | 0 |
| Other neurodegenerative disorders | 0.4 | 1 | 0 |
| Gastrointestinal problems | 6 | 5 | 7 |
| Thyroid problems | 2 | 3 | 1 |
| Cancer | 3 | 2 | 3 |

## Results

### Shotgun metagenomics reveals a defined aging-associated shift in microbial richness and functional guilds

To study the role of the gut microbiome in healthy aging, stool samples from a deeply phenotyped cohort of community-living Asian octogenarians (SG90, **Table 1**, **Supplementary Data File 1**) were analyzed with deep shotgun metagenomic sequencing (n=234, 17 million reads on average, **Methods**). The resulting high-resolution species-level taxonomic profiles were compared to reference shotgun metagenomic data from healthy, younger Singaporeans^34, 35^ from two cohorts (SPMP, n=109, age range=[53-74]; CPE, n=96, age range=[21-80]), as well as other Asian populations^36^ (T2D, n=171, age range=[21-70]; **Supplementary Data File 2**). Systematic joint analysis of these cohorts after batch-correction and matching other demographic characteristics (gender, ethnicity; **Methods**), revealed a progressive shift in taxonomic profiles across different age groups (n=516, age range=[21-100], **Figure 1A**), particularly along the y=x axis relative to the first and second principal components of variation (ordinal logistic test p-value<0.001, **Figure 1B, Supplementary Figure 1A**). Diversity analysis indicated that this was accompanied by an aging-associated shift (ordinal logistic test p-value<0.001, **Figure 1C**) that is driven by microbial richness and evenness, both of which exhibit significant reduction with age (**Supplementary Figure 1B-D**). PERMANOVA analysis showed that Age was the major source of variation compared to other attributes (**Supplementary Figure 2A**) and that beta diversity peaked first in the 41-60 age group and then again in the 91-100 age group (**Supplementary Figure 2B**), though uniqueness did not consistently increase with age as reported previously in a western cohort^27^. These results highlight an aging-associated shift in microbial composition that is defined by the loss of gut species, and increase in beta diversity in extreme age groups.

**Figure 1:**
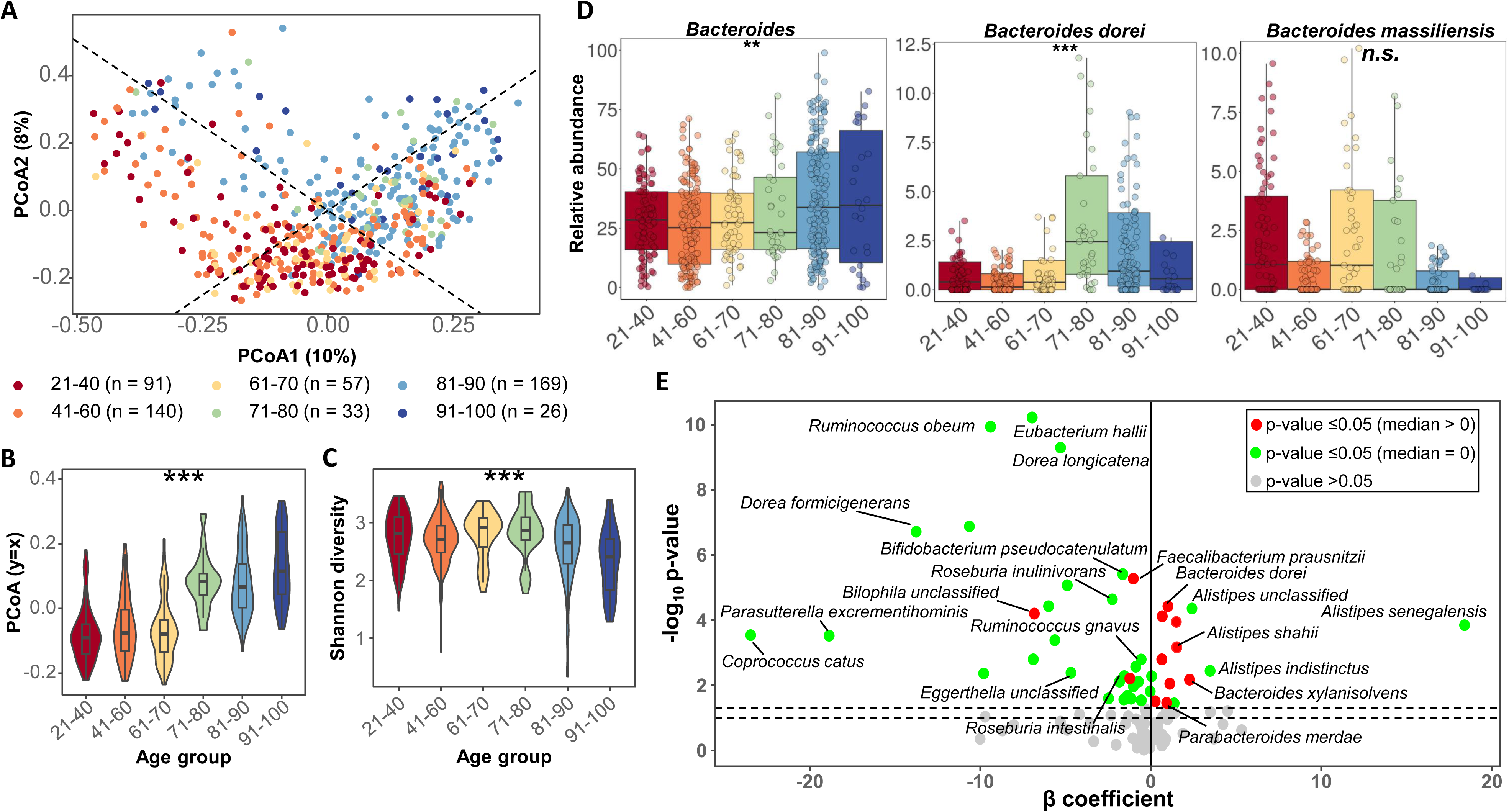
Aging-associated shifts in gut microbiome richness and relative abundance of key species. (A) Principal coordinates analysis (PCoA) plot based on species-level Bray-Curtis dissimilarity of gut microbiome profiles across age groups after batch correction. Dashed lines indicate the y=x and y=-x axes. (B-C) Violin plots showing the distribution across age groups of (B) PCoA1 and PCoA2 values projected on the y=x axes and (C) Shannon diversity indices. The symbols ‘**’ and ‘***’ represent p-value<0.05 and p-value*<*0.005 (Ordinal logistic test), respectively. (D) Relative abundance boxplots for the *Bacteroides* genus, and corresponding species that was identified to be associated significantly and not significantly with age. The symbols “*n.s.”*, “*” and “***” represent p-value>0.05, p-value≤0.05 and p-value<0.005, respectively (FDR-adjusted; GLM test for association with age). (E) Volcano plot showing the β coefficient on the x-axis and FDR-adjusted –log_10_ p-values (GLM test for taxa association with age) on the y-axis. Points for statistically significant taxa from “high stringency list” and those with high beta coefficients are highlighted in the volcano plot as red colored points for median abundance in the SG90 cohort greater than zero and green otherwise, with non-significant taxa shown with grey dots.

In order to identify taxa associated with age, a generalized linear model (GLM) was used to account for demographic and clinical covariates that can be confounders (e.g. gender, body mass index, fasting blood glucose, triglyceride, total cholesterol and high density lipoprotein; **Methods**). Overall, 4 phyla, 22 genera and 40 species were found to be associated significantly with age (FDR-adjusted p-value<0.05, **Supplementary Table 1**, **Supplementary Data File 3**). The phylum and genus level results were broadly consistent with prior observations (e.g. in Chinese^37^, Japanese^25^ and Italian centenarians^28, 38^), with enrichment in Bacteroidetes (β=0.33, FDR-adjusted p-value<0.001), and depletion in Firmicutes (β=-0.49, FDR-adjusted p-value<0.001) with age (**Supplementary Table 1**). At the genus level a strong enrichment in unclassified *Acidaminococcaceae* species (β=6.69, FDR-adjusted p-value<0.05) and depletion in *Parasutterella* species (β=-20.19, FDR-adjusted p-value<0.001) was observed that has not been noted before, while at the species level a majority of our observations were novel owing to the higher resolution of shotgun-metagenomics^20–26^ (**Supplementary Data File 3**, **Table 2**). In addition, we tested the robustness of this analysis by varying the normalization approach, statistical model and cohorts used to find that most taxa associations were detected in a majority of the conditions tested (**Methods**, **Supplementary Figure 3**). We also noted that while several species associations were detected in individual cohorts (n=10), the inclusion of the SG90 cohort notably boosted the number of age-associations identified (n=38), suggesting that having a wider age range and more subjects could benefit such analysis (**Supplementary Figure 4**). To mitigate the potential effect of incomplete batch-correction, a high-stringency list of species associations was further derived based primarily on methods that do not depend on this procedure (n=17; **Table 2**, **Figure 1D**-**E**, **Supplementary Figure 3**; **Methods**). Reanalysis of taxonomic profiles and inclusion of results from other publicly available cohorts^17, 39–43^ (n=6; **Methods**) highlighted the consistency of these associations with a high percentage being replicated or validated (27/40=68%, odds ratio=21.4, p-value<6×10^-15^). In addition, the overlap with Asian cohorts (18/40, odds ratio=8.83, p-value<10^-6^) was more statistically significant than with western cohorts (16/40, odds ratio=3.10, p-value<7.1×10^-3^), despite having more western cohorts in this comparison (4/6), highlighting the value of having more Asian datasets focused on healthy aging (**Supplementary Figure 5**).

**Table 2:** Species-level gut microbial taxa associated with aging. . The table reports statistically significant taxa based on FDR-adjusted p-value≤0.05 with a GLM test and covariate adjustment. Species with median relative abundance greater than zero in the SG90 cohort are reported here and the full list is provided in **Supplementary Data File 3**. The β coefficient estimates strength of association as change in age (in years) for every 1% increment of taxa relative abundance. Taxa in the high-stringency list are shown in bold face.

| Enriched with age | p-value | $\beta$ coefficient for age<br>[95% C.I.] | Remarks |
| --- | --- | --- | --- |
| ***Bacteroides xylanisolvens*** | $6.71 \times 10^{-3}$ | 2.28 [1.05, 3.51] | |
| ***Alistipes shahii*** | $6.81 \times 10^{-4}$ | 1.53 [0.99, 2.07] | |
| *Bacteroidales bacterium ph8* | $1.13 \times 10^{-4}$ | 1.51 [0.83, 2.19] | |
| *Klebsiella pneumoniae* | $8.92 \times 10^{-3}$ | 1.12 [0.39, 1.86] | |
| *Escherichia unclassified* | $3.73 \times 10^{-5}$ | 1.02 [0.59, 1.45] | 16S OTU enriched in Indian centenarians<sup>150</sup> |
| ***Parabacteroides merdae*** | $3.42 \times 10^{-2}$ | 0.95 [-0.0087, 1.90] | |
| ***Bacteroides dorei*** | $7.61 \times 10^{-5}$ | 0.68 [0.42, 0.94] | |
| *Alistipes onderdonkii* | $1.60 \times 10^{-3}$ | 0.66 [0.30, 1.02] | Enriched in Italian centenarians<sup>28, 151</sup> |
| *Escherichia coli* | $3.10 \times 10^{-2}$ | 0.27 [0.054, 0.48] | |
| **Depleted with age** |  |  |  |
| ***Bilophila unclassified*** | $6.30 \times 10^{-5}$ | -6.82 [-9.90, -3.74] | |
| *Ruminococcus torques* | $6.14 \times 10^{-3}$ | -1.22 [-1.99, -0.45] | |
| ***Faecalibacterium prausnitzii*** | $5.33 \times 10^{-6}$ | -1.01 [-1.29, -0.74] | Depleted in Italian centenarians<sup>28, 108</sup> |

Among the core gut microbiome taxa identified^27, 44^, only some species in the *Bacteroides* genus showed significant positive associations including *Bacteroides dorei* and *Bacteroides xylanisolvens*, with *Bacteroides massiliensis* even exhibiting a slight negative trend, though this variation is masked at the genus level highlighting the utility of species-level resolution offered by shotgun metagenomics (**Table 2**, **Figure 1D-E**). We also noted an enrichment with age for multiple *Alistipes* species, including *Alistipes shahii, Alistipes onderdonkii* and *Alistipes senegalensis*, that are bile tolerant, have a unique way of fermenting amino acids such as L-lysine^45, 46^, and can produce neurotransmitter precursors such as indole^47^ (**Supplementary Figure 6A**). In contrast, several species that are known to be important for butyrate production in the gut^45, 46^, including *Faecalibacterium prausnitzii*, *Roseburia inulinivorans* and *Eubacterium hallii*, were significantly depleted with age (**Figure 1E**, **Supplementary Figure 6B**). Several of these associations were seen in other Asian cohorts but not in the western cohorts analyzed here (n=11, e.g. *Dorea formicigenerans*, *R. inulinivorans*, *A. shahii* and *Eubacterium hallii*) indicating that there may be consistent key differences in ethnic Asian populations. Also, in a few cases the directionality of association differed between Asian and western cohorts (e.g. *B. xylanisolvens*), highlighting the importance of population-specific studies. Integrating this information through taxonomic co-occurrence analysis highlighted a distinct network structure in the elderly (32 edges and 3 hubs) compared to younger individuals (80 edges and 9 hubs), with a shared pattern of cliques formed by the classical butyrate producers and *Alistipes* species, indicating that these may represent distinct functional *guilds*^48^ that switch roles with age (**Supplementary Figure 7**).

### Gut microbiomes in elderly Asians exhibit a metabolic potential expansion in butyrate synthesis pathways

To further analyze metabolic changes associated with aging, shotgun metagenomic data from all subjects was used to quantify gene and pathway abundances (**Supplementary Data File 4**, **Methods**). In total, 413 metabolic pathways were identified and quantified of which 70 were found to be differentially abundant across various age groups (**Supplementary Data File 5**), broadly grouping into the four categories of sugar metabolism (17 pathways), vitamin, energy metabolism and cell wall biosynthesis (28 pathways), lipid metabolism (9 pathways) and amino acid metabolism (16 pathways, **Figure 2**). Specifically, among sugar metabolism pathways we observed distinct age-associated biases, where e.g., pyruvate fermentation to acetate and lactate was enriched in the younger age group (21-40), while degradation of mono-, di- and polysaccharides was enriched in older age groups (**Figure 2A**). As many as 8 different pathways for various substrates were enriched in the 61-70 age group, including simple pentose sugars (Pentose phosphate pathways), and complex polysaccharides such as stachyose and starch (Starch degradation V). Notably, microbial glycolytic pathways for metabolism of simple sugars (glucose) were enriched in healthy individuals over 90, while prior work had highlighted the role of corresponding pathways in aging in model organisms^49^.

**Figure 2:**
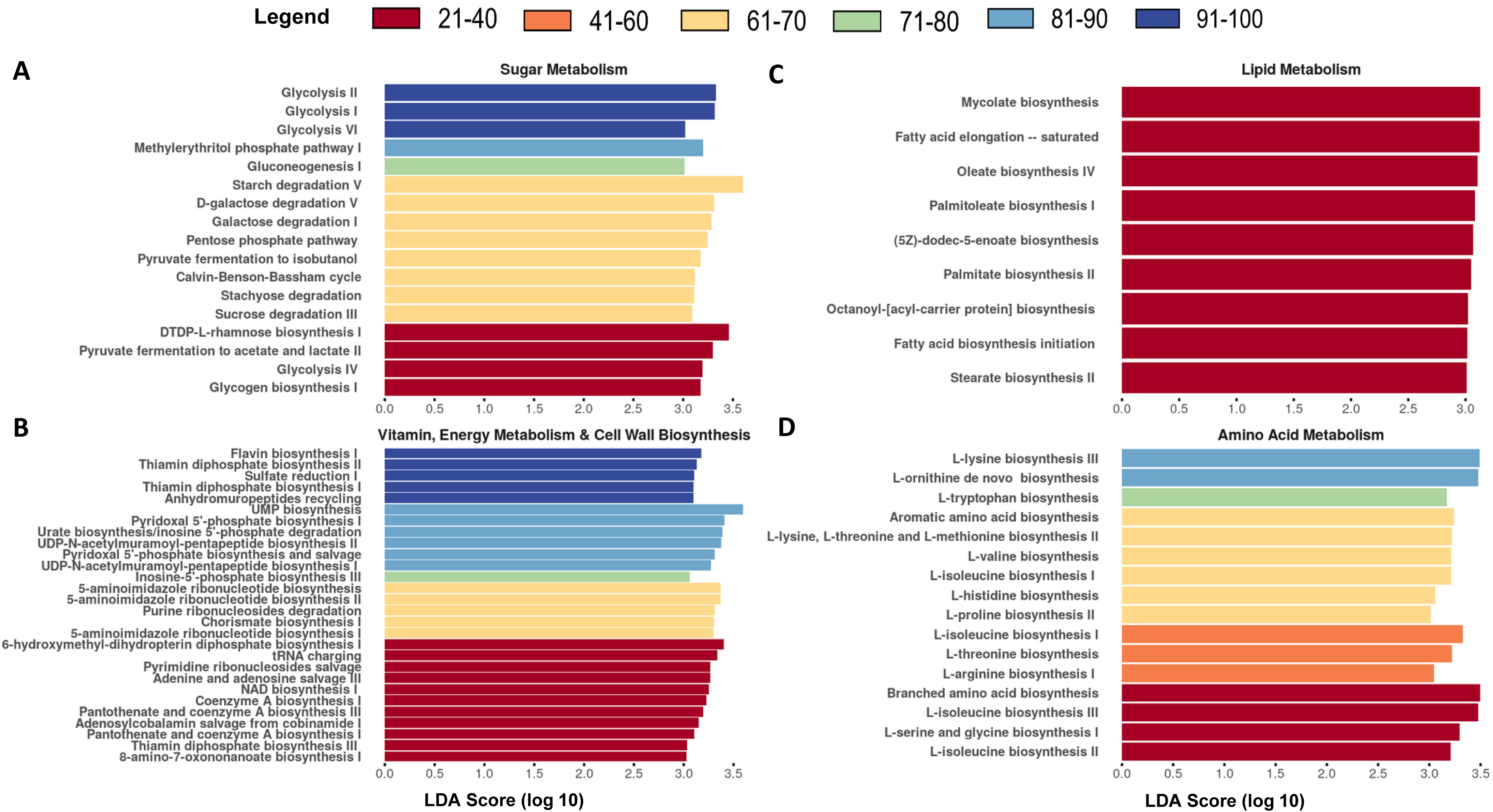
Associations of microbial metabolic pathways across age groups. Barplot showing pathways that are significantly associated and enriched specifically in different age groups (p-value≤0.05) based on LEfSe analysis. Pathways were grouped into the broad categories of (A) Sugar metabolism, (B) Vitamin, energy metabolism and cell wall biosynthesis, (C) Lipid metabolism and (D) Amino acid metabolism. The effect size is depicted in the form of an LDA Score (log 10).

In line with the vital role of vitamins as essential micronutrients (particularly in energy metabolism and immune function^50, 51^) with limited synthesis in humans^50^, several microbial pathways were associated with aging (especially B vitamins, **Figure 2B**). In particular, thiamine diphosphate and flavin biosynthesis were enriched in the most elderly (91-100) and could have anti-inflammatory and anti-aging roles^51^. On the other hand, three alternate biosynthetic pathways to produce thiamin and pantothenate were enriched in younger individuals (21-40).

While microbial lipid metabolism pathways were predominant in younger subjects (21-40 age group, **Figure 2C**), pathways related to amino acid metabolism were more often enriched in the elderly (particular 61-70 age group), including the synthesis of various essential amino acids such as histidine, valine, threonine, isoleucine, tryptophan, and methionine and aromatic amino acids (**Figure 2D**). In addition, lysine biosynthesis was over-represented in the gut microbiomes of two older age groups^52^ (61-70 and 81-90), linking to our earlier observation of age-related enrichment in specific *Bacteroides* and *Alistipes* species, which are known to synthesize lysine and produce short-chain fatty acids such as butyrate with lysine as substrate^45^, respectively.

Analysis of the four major known pathways for butyrate production^45, 46^ (from pyruvate, glutarate, 4-amino butyrate and lysine) confirmed that while several lysine-associated genes show an age-related increasing trend (e.g. *kamA,* FDR-adjusted p-value<0.01 and *kdd,* FDR-adjusted p-value<0.05), no significant associations were seen in the pyruvate related genes (**Figure 3**). Furthermore, the overall gene abundance in the pathway for butyrate production from lysine also showed a significant increase with age (FDR-adjusted p-value<0.001, **Supplementary Figure 8**). Metabolic pathways analysis revealed that the abundance of gut microbial species that provide metabolic support to butyrate producers peaked in older age groups (71-80; **Supplementary Figure 9**, **Methods**). To more directly link taxonomic and functional analysis, we first analysed the contribution of taxa to L-lysine biosynthesis III pathway and observed that *Bacteroides sp.* were the dominant contributors, with the contribution of *B. xylanisolvens* increasing with age (**Supplementary Figure 10A, 10B).** Next, we systematically generated metagenome-assembled genomes, annotated genes and pathways, and conducted metabolic network analysis (**Methods**). This analysis showed that the metagenome-assembled genomes represented a diversity of *Bacteroides* and *Alistipes* species, with many genomes harboring corresponding complete pathways for lysine biosynthesis (*Bacteroides*) and lysine-to-butyrate conversion (*Alistipes*) that are reachable based on metabolic network analysis (**Supplementary Figure 10C-E**). These observations, along with those in the previous section, indicate a metabolic potential expansion in butyrate synthesis pathways from precursors other than pyruvate in the gut microbiomes of elderly Asians.

**Figure 3:**
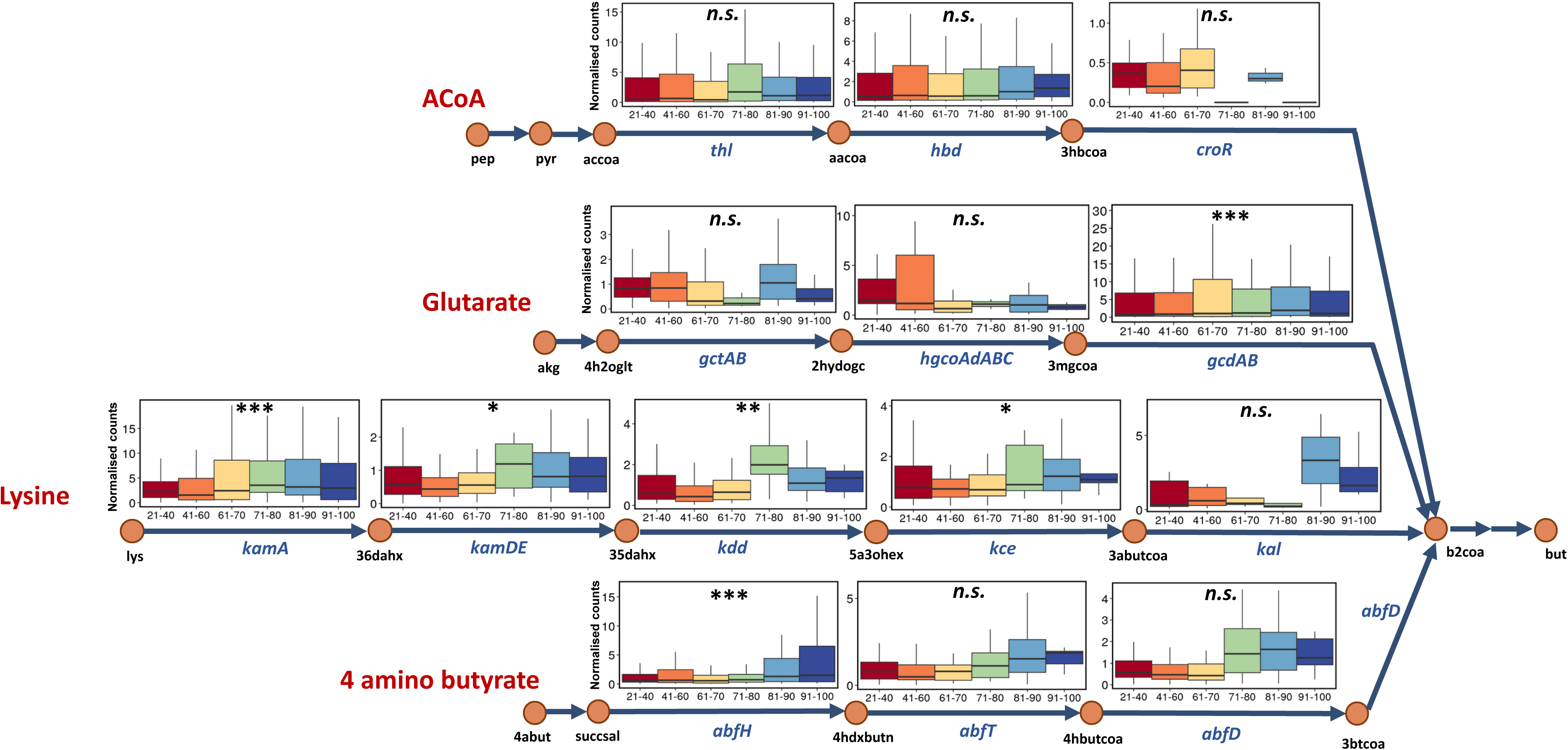
Differential representation of butyrate synthesis pathways in gut metagenomes across age groups. Pathway diagram where nodes represent major metabolites involved in various steps of butyrate production (but) starting from 4 different precursors (pyruvate, glutarate, lysine and 4-aminobutyrate), and edges are labelled with the genes involved in the conversion. Boxplots present normalized counts for the corresponding gene across different age groups. The labels “*n.s.”,* “*”, “**” and “***” represent p-value>0.05, p-value≤0.05, p-value<0.01 and p-value<0.001, respectively (FDR-adjusted GLM test for gene association with age).

To further explore these findings, we leveraged the recent development of a healthy aging model in mice, where dietary supplementation with alpha ketoglutarate (AKG) to aged mice (18 months) not only increased lifespan but also improved inflammation and frailty markers^53^. Shotgun metagenomics of fecal samples from AKG-supplemented and control mice (n=20 per group, **Supplementary Figure 11A**; **Methods**) showed that even though mice harbor distinct gut microbial species, at the pathway level a similar expansion in butyrate production pathways is seen in healthy aging mice (**Supplementary Figure 11B**). In particular, the lysine to butyrate conversion pathways shows the strongest enrichment, as seen in our human data (1.65-fold increase in median abundance, FDR-adjusted p-value<0.01). At the gene level, several key genes for lysine to butyrate production were strongly enriched including *kdd*, *kamDE*, *kce* and *kamA* (FDR-adjusted p-value<0.05; **Supplementary Figure 11C**). Such a trend was not observed for genes involved in pyruvate to butyrate conversion, highlighting the role of expansion in alternate butyrate synthesis pathways to support healthy aging.

### Robust species-level associations between gut microbial taxa and aging phenotypes

Aging is a significant risk factor for chronic diseases impacting multiple organ functions, including metabolic, immune, and musculoskeletal systems, which could be mediated via gut microbiome function^26, 54^. To further relate the taxonomic and functional changes observed in Asian octogenarians with their clinical phenotypes we harnessed the higher resolution of shotgun metagenomic data, broader age range and size of our cohorts, to robustly identify associations in a generalized linear model framework while accounting for demographic and clinical confounders (e.g. age, gender, other metabolic markers; **Methods**, **Supplementary Data File 6**). Despite the multiplicity of taxa and clinical features, we noted that this analysis typically resulted in a few significant associations (**Figure 4**). For example, for fasting blood glucose as a well-studied biomarker for glucose metabolism^55^, insulin resistance^56^, and type 2 diabetes^57^, we identified two associated species (*Bifidobacterium adolescentis* [β=0.08, FDR-adjusted p-value<0.01] and *Parabacteroides goldsteinii* [β=0.48, FDR-adjusted p-value<0.01], **Figure 4A**) with prior *in vitro* and *in vivo* observations supporting this link^58–60^, as well as a new association with a *Lactobacillus* species (*Lactobacillus mucosae:* β=0.88, FDR-adjusted p-value<0.01) that has never been described before despite numerous studies for this genus^61–64^. Note that β values indicate the change in response variable (e.g. fasting blood glucose levels) for every unit change in relative abundance values, after accounting for other confounding variables. Given the known normal range of fasting blood glucose levels (<6.1 mmol/L^65^; **Supplementary Table 2**) the observed β values (0.48 and 0.88) therefore could suggest a large effect size.

**Figure 4:**
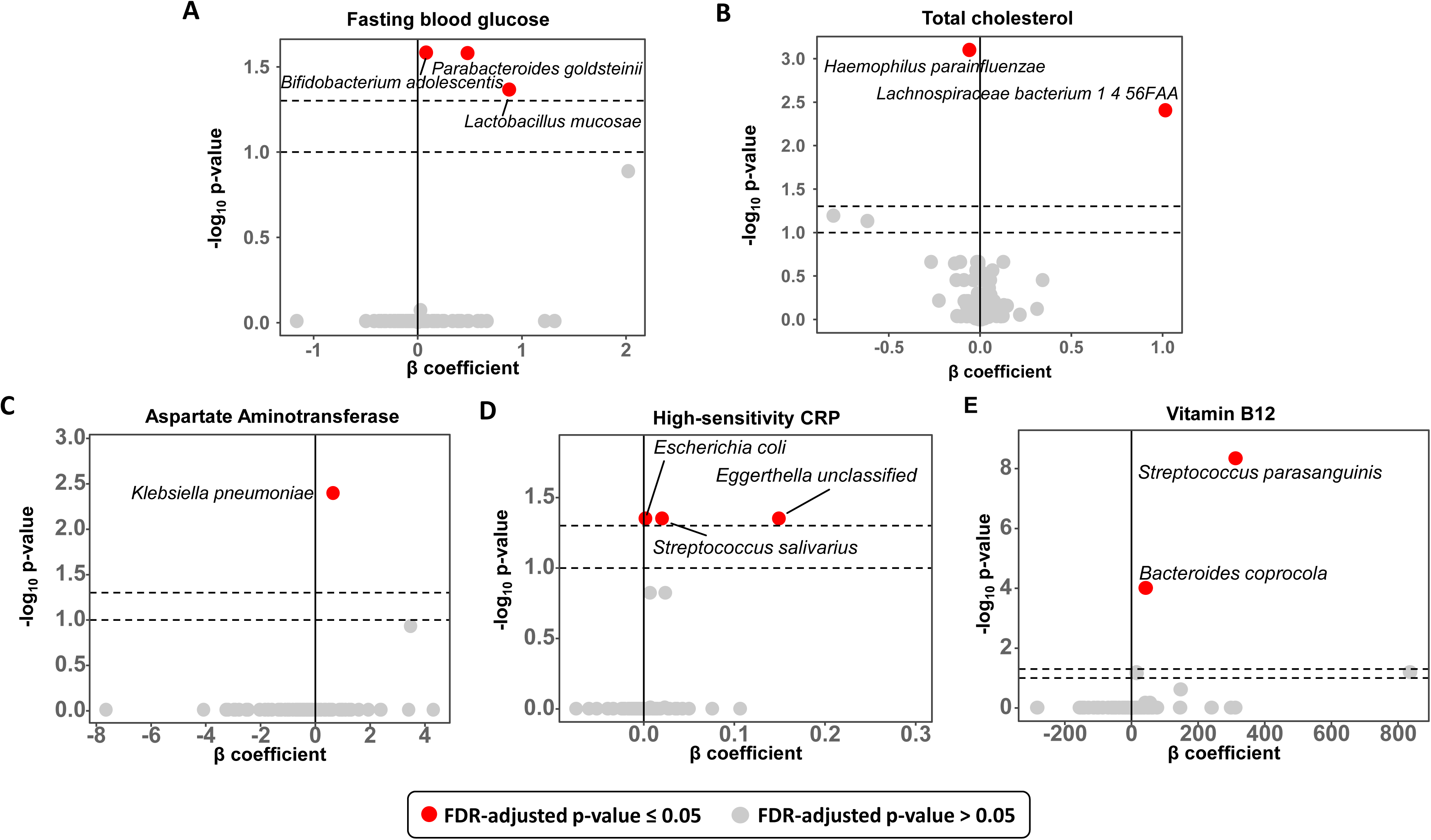
Microbiome associations with key clinical markers. Volcano plots showing the β coefficient on the x-axis and the FDR-adjusted –log_10_ p-values on the y-axis based on GLM test for gut microbial associations with clinical markers for (A) Glucose control, as measured by fasting blood glucose levels, (B) Lipid control, as measured by total cholesterol in plasma, (C) Liver health, as measured by plasma levels of aspartate aminotransferase, and (D) Systemic inflammation, as measured by high-sensitivity CRP levels, (E) Vitamin B12. Points in all plots are colored with red for p-value≤0.05 and grey otherwise. Dotted lines represent the thresholds of p-value*=*0.05 and p-value*=*0.1.

We next analyzed cholesterol biomarkers (e.g. high-density lipoproteins - HDL, low-density lipoproteins – LDL and triglycerides) for hypercholesterolemia and cardiovascular diseases, due to their impact on healthy aging^66^. Association analysis with total cholesterol levels revealed two significant candidates, positive association with abundance of *Lachnospiraceae bacterium 1 4 56FAA* (β=1.02, FDR-adjusted p-value=3.92×10^-4^) and negative association with *Haemophilus parainfluenzae* (β=-0.06, FDR-adjusted p-value=7.95×10^-5^, **Figure 4B**), neither of which has been reported previously^67, 68^. Interestingly, analysis with LDL levels revealed the opposite trend, with the *Lachnospiraceae* species (β=-0.88, FDR-adjusted p-value=4.27×10^-2^) being negatively correlated and *H. parainfluenzae* (β=0.06 FDR-adjusted p-value=1.72×10^-3^) being positively correlated, and similar trends being observed with HDL but not reaching statistical significance (**Supplementary Figure 12A, 12B**). These observations indicate potential beneficial roles of *Lachnospiraceae* species in maintaining lower levels of LDL. Interestingly, microbial association with triglyceride levels was dominated again by positive association with *Haemophilus parainfluenzae* (**Supplementary Figure 12C**), a species that has been described to be strongly enriched in patients with chronic cholecystitis patients^69, 70^ and individuals with increased risk of cardiovascular disease^71, 72^.

As the biological processes of aging are in part believed to be accelerated by inflammation^73,74^ (*Inflammaging*), including risk for cardiovascular disease, frailty and disability^75–77^, we sought to identify gut microbial associations with inflammation markers. Specifically, serum levels of aspartate aminotransferase (AST) were associated with the abundance of *Klebsiella pneumoniae* (β=0.65, FDR-adjusted p-value=3.99×10^-3^, **Figure 4C**), a sporadic colonizer of the gut microbiome, and a known pathogen for liver abscesses that are increasingly common in Asian populations^78^. A similar association was seen with alanine aminotransferase (ALT), a more specific biomarker for liver injury, but it did not reach statistical significance (**Supplementary Figure 12D**). Analysis with high-sensitivity C-reactive protein (hs-CRP) levels, a commonly used marker for systemic inflammation^79^, identified a known association with *Escherichia coli*^80^ (β=0.0020, FDR-adjusted p-value=0.04), as well as an association with a *Streptococcus* species (*S. salivarius*: β=0.02, FDR-adjusted p-value=0.04) which has been linked previously with rheumatoid arthritis^81^ (**Figure 4D**). Furthermore, a significant association was observed with a *Eggerthella* species (β=0.15, FDR-adjusted p-value=0.04) distinct from *Eggerthella* lenta^82^, whose role is less well-understood^67^, and the association with host inflammation has not been described before. These results highlight the various bacterial species and potential pathogenic functions by which altered gut microbiomes can contribute to inflammaging.

Aging is a common risk factor that often predisposes elderly to the development of neurological disorders such as Parkinson’s and Alzheimer’s disease^83^. Many studies have shown that Vitamin B12 is involved in several processes of the nervous system, including the synthesis of myelin and post-injury nerve regeneration^84, 85^, the deficiency of which causes peripheral neuropathy^86^, gait ataxia and physical frailty that are often observed in the elderly. A strong association was observed between serum levels of Vitamin B12 and the bacterial species *Streptococcus parasanguinis* (β=312.50, FDR-adjusted p-value<5×10^-9^, **Figure 4E**), strains of which are known to produce Vitamin B12^87^, highlighting the metabolic potential for gut bacteria to support healthy aging. Intriguingly, we also observed a significant association for blood levels of Vitamin B12 and the abundance of *Bacteroides coprocola* (β=42.89, FDR-adjusted p-value<10^-4^), even though Bacteroidetes are not common B12 producers^88^ and instead compete with the host to capture B12 in the gut^89^, indicating that dietary intake might be an explanation for the observed positive association. Together, these observations highlight the two-way communication between gut microbial species and their environment that can impact host phenotypes as well as serve as meaningful biomarkers of healthy aging.

## Discussion

The complexity of the human gut microbiome, both in terms of its genetic diversity and the myriad host-microbial interactions that shape it, necessitates that association studies be done with sufficiently high-resolution and power to help infer meaningful insights about function and mechanisms^90^. While prior studies were primarily limited by their resolution (16S rRNA surveys^20–26^), recent studies have underscored the utility of shotgun metagenomics for species-level and functional pathway analysis, albeit in limited-sized cohorts^28, 29^. By generating deep shotgun metagenomic data for >200 elderly subjects, this study significantly enhances our ability to explore the role of the microbiome in healthy aging. In particular, we provide the first well-powered species-level view of microbiome shifts as a function of age, with a steady decrease in species richness (**Figure 1A**, **Supplementary Figure 1B**). This is accompanied by an increase in beta diversity in extreme age groups, but further defines that this is due to an overall shift of microbial community composition with age, with specific species serving as markers (e.g. *A. shahii*, *B. xylanisolvens, R. obeum* and *E. hallii*). Our analysis also highlights that the age-associated shift is on an axis of variation (**Figure 1A-B)** that is distinct from the axis separating different gut microbiome clusters reminiscent of enterotype structures^91^ (**Supplementary Figure 1A**).

In addition, while the overall reduction in microbial diversity observed here (**Figure 1C**) is broadly consistent with some studies^24, 92^, our analysis clarifies that this is accompanied by an increase in beta diversity of microbiome composition and a loss of species richness (**Supplementary Figure 2B, 1B-D**), defined in part by several classical butyrate producers in the gut (e.g. *F. prausnitzii*, *R. inulinivorans* and *E. rectale*). We did not observe a general loss in core *Bacteroides* species as reported in Wilmanski et al^27^, which could be a function of our Asian cohorts, but may also be due to methodological differences (e.g. covariate adjustment). Also, our results suggest that the trends reported for richness^22, 93^ and uniqueness^27^ may be variable across cohorts^94^. Further studies are needed across different global populations to explore the factors impacting this variability using consistently generated datasets and potentially using long read sequencing to enable more reliable detection/assembly of rare taxa. This could also impact the detection of associations with viruses and fungi (none reported here), which are only sporadically abundant in the human gut microbiome.

In order to have data from a wide-range of ages and a large number of subjects to sufficiently power our age-association analysis, we combined data from multiple Asian cohorts (SG90, T2D, SPMP, CPE) with metagenomic sequencing data, and attempted to account for differences in how the data was generated using a batch-correction approach (**Methods**, **Supplementary Table 4**). In addition, our association analysis accounted for several demographic and clinical factors that could be confounders, including ethnicity, gender, body mass index, fasting blood glucose, triglyceride, total cholesterol and high density lipoprotein levels lipoprotein. Nevertheless, it is important to note that batch-correction methods cannot be expected to fully account for all technical sources of variation, and similarly, other potential confounders (e.g. diet, medications) that could not be accounted for as this data was not consistently available across cohorts. Multi-cohort association analysis thus necessarily has to evaluate the robustness of the results obtained as a function of various data subsets used, technical choices made (e.g. normalization) and statistical approaches used. Our experiments suggest that the batch-correction and GLM-based association analysis reported here provide robust results that are reproduced across a majority of conditions tested (**Methods**, **Supplementary Figure 3**). This was further validated by the strong enrichment of age associations detected in other cohorts, particularly in Asian cohorts (**Supplementary Figure 5**). In addition, our analysis highlights the importance of having sufficient data from older age groups (**Supplementary Figure 4-5**). Overall, despite all analytical care, the interpretation of microbiome associations with aging needs care as it is a multi-factorial phenomenon, where e.g. increased lifetime use of antibiotics could explain loss in gut microbial species richness with age, while at the same time such observations could arise from other factors such as lifestyle choices (e.g. smoking) and their independent impact on longevity and the gut microbiome. Our study is structured to identify microbiome associations with age and considers BMI, total cholesterol, fasting blood glucose etc. as confounders, though in principle, the influence of age on these variables could be through the microbiome. In that sense, our study is conservative and should identify only the strongest independent associations. While we would ideally identify causal associations between the microbiome and healthy aging, bidirectional communication between the host and the microbiome makes this interpretation challenging from a cross-sectional study. Nevertheless, on their own, the associations identified here can serve as meaningful biomarkers that allow us to track healthy aging. In combination with more controlled intervention studies, such as the mouse model of healthy aging studied here, we can then refine hypotheses on microbial pathways that play a causal role in healthy aging.

The importance of incorporating age and other covariates in microbiome association analyses has been emphasized in several recent studies^17, 95^. While this also necessitates larger cohort sizes and extensive metadata collection, a potential advantage is that the few associations that survive covariate adjustments are likely to be more meaningful (e.g. no associations were detected for sleep duration, frailty markers or free thyroxine levels; **Supplementary Data File 6**). In combination with shotgun metagenomics, where specific species and functional genes/pathways can be directly quantified, and related through metagenome-assembled genomes, this provides for a powerful avenue for developing meaningful mechanistic hypotheses informed by *in vivo* human data. In particular, a striking feature of our results is the depletion of a ‘microbial guild’ of butyrate producing species in elderly subjects and its replacement with specific *Alistipes* species that can produce butyrate through an alternate pathway using amino acids as precursors. At the metabolic potential level, we do not see a significant reduction for the pyruvate-to-butyrate conversion pathway, and it is not clear if dietary or microbially produced amino acids are the key source for the alternate pathways^96^. Prior studies have shown reduction in short-chain fatty acid (SCFA) levels, particularly butyrate, in stool samples from the elderly^97^, association between butyrate levels and frailty^14^, and hypothesized a shift towards bacterial protein fermentation due to increased intestinal transit time^98^. We observed a relative reduction of microbial to host reads in elderly subjects (**Supplementary Figure 13**) that could reflect a reduced capability to support gut microbial fermentation, though direct measurements are needed to confirm this indirect evidence. Changes in dietary patterns (e.g. low fiber) as well as intestinal physiology (e.g. muscle loss, hypochlorhydria) could thus partly explain the observed shift in species composition and deserves further investigation. Our data from a mouse model of healthy aging further supports the idea that alternate butyrate production pathways in the gut microbiome could be important with age. The model is established by feeding mice α-ketoglutarate which is rapidly absorbed in the small intestine^99^, but may still explain the enrichment seen in the glutarate pathway in the gut microbiome. The strong enrichment seen in the lysine-to-butyrate pathways is consistent with our observations in human cohorts and could reflect the systemic metabolic and physiological effects of α-ketoglutarate in promoting healthy aging^100^, though it is not clear if this effect could be mediated through the gut microbiome. Follow-up experiments in mice with defined microbiome compositions, with and without α-ketoglutarate supplementation, could help further disentangle these relationships.

As the microbial production of SCFAs in the gut is known to play an important role in many aspects of human health^101, 102^ (e.g. gut barrier function, immune regulation, gut-brain communication), it is not surprising therefore that reduction in butyrate-producing species has been associated with frailty^103^. Based on the observations in this study we hypothesize that alternate pathways for butyrate production, including using glutarate, 4-amino butyrate and lysine as precursors, may partially augment the pyruvate-dependent pathway in the gut microbiomes of elderly individuals to promote healthy aging. These observations may be due to a “selection on survival” bias^27^, but results from the mouse model suggest that this may not provide a full explanation. Lysine is an essential amino acid that is found in lower plasma concentrations in frail elderly subjects^104^, and lysine supplementation is being investigated in multiple clinical trials for its health benefits. Its use as a supplement to promote healthy aging could thus be another avenue to explore via the pathway of boosting butyrate production in the gut. Among other pathways that were relatively depleted in the elderly, reduction in pyruvate fermentation to acetate and lactate and lipid metabolism pathways (**Figure 2**) may have detrimental effects for host health and can be targets for compensation through supplements and probiotics^105, 106^. On the other hand, the relative enrichment of vitamin synthesis pathways in the elderly, including thiamine diphosphate (vitamin B_1_), flavin (vitamin B_2_) and phospho-pantothenate (vitamin B_5_), as well as the association of specific species with serum vitamin B_12_ levels (**Figure 4E**), could contribute to healthy aging and thus deserves further study.

Several aging-related chronic diseases, including type 2 diabetes^36^, cardiovascular diseases^107^, and neurological conditions^108^ have been linked with the gut microbiome and thus it is essential to account for age as a confounding factor when studying gut microbial mechanisms that contribute to disease. We hypothesize that having individuals from as wide a range of ages as possible, particularly those who have aged well, will likely help strengthen and refine such association analysis. We detected a positive association of *Bifidobacterium adolescentis* with fasting blood glucose, which is well-known to impact glycemic control^109^. In addition, we also noted a significant association for *P. goldsteinii*, a potential new probiotic species with emerging evidence for a diet-dependent role in obesity and type 2 diabetes^58, 110^, though the mechanisms for this effect in mouse models remain to be elucidated.

Systemic low-levels of inflammation are believed to contribute to various aging-related phenotypes^73, 74^ (*inflammaging*). Correspondingly, associations between gut microbial species and markers of inflammation (e.g. hs-CRP and AST) are of particular interest. While the association of a common-source of liver abscesses in Asia (*K. pneumoniae*) with a liver disease marker (AST), and a common oral bacterium (*S. salivarius*) with a systemic inflammation marker (hs-CRP) are consistent with disease biology, the distinctness of these associations is striking i.e., while *K. pneumoniae* is the only species associated with AST, a more diverse group of bacteria are associated with hs-CRP levels. Furthermore, relatively little is known about the *Lachnospiraceae* species with strong associations to cholesterol/LDL levels. Together with the strong association of microbial biomarkers such as *S. parasanguinis and B. coprocola* with serum Vitamin B_12_ levels, these findings could help develop a non-invasive frailty test based on at-home sample collection^111^. Overall, these results highlight the value of species-level shotgun metagenomic analysis in large well-characterized cohorts, where further studies in other populations would help determine if some of these associations are specific to Asian environments and lifestyles.

## Methods

### Cohorts and datasets

#### SG90 cohort

The SG90 cohort is based on a longitudinal population health study that was set up in the 1990s which involved routine measurement of metabolic and other health variables^30^. The current dataset is based on a subset of 234 elderly individuals (77-97 years old) who are community-living participants (not living in a nursing home, no diagnosis of dementia and not physically unfit) and consented to providing their stool and blood samples. The participants in the SG90 cohort were recruited under the SLAS-3^112^ protocol approved by the Institutional Review Board (IRB) at National University of Singapore (reference number: B-15-081). This study was also approved by an IRB for the Singapore Chinese Health Study^30^ (reference number: H-17-027, sample collection: 2019/00439). Fasting blood glucose (mmol/L), triglyceride (mmol/L), total cholesterol (mmol/L), HDL (mmol/L), LDL (mmol/L), hs-CRP (mg/L), AST (U/L), ALT (U/L) and Vitamin B12 levels (pmol/L) were measured based on a blood draw. Stool samples were collected either at the time of blood collection or within a week. In addition, physical assessments were performed to calculate BMI (kg/m^2^) from their weight (kg) and height (m)., as well as measure gait speed (m/s) and handgrip strength (kg) of subjects.

#### SPMP dataset

The SPMP dataset is based on the recall of a subset of 109 healthy Singaporean subjects (53-74 years old) from a multi-omics study in Singapore^113^. Stool samples were collected for gut microbiome analysis using shotgun metagenomic sequencing^34^.

#### T2D dataset

Shotgun metagenomic datasets were obtained for 171 healthy Chinese individuals from a previously published type 2 diabetes (T2D) study^36, 114^ using the curatedMetagenomicData package^114^. Briefly, the subjects chosen for our study were 21-70 years old and were non-diabetic controls in the study. Clinical data such as fasting blood glucose (mmol/L), triglyceride (mmol/L), total cholesterol (mmol/L), HDL (mmol/L) and LDL (mmol/L) levels were also obtained from this study^36^.

#### CPE dataset

The CPE dataset^35^ is based on a prospective cohort study consisting of CPE-colonized subjects and their healthy family members. For our comparisons, we used shotgun metagenomic data of 82 healthy family members (21-80 years old) with Chinese ethnicity.

#### Demographic matching

Analysis for all cohorts was restricted to ethnic Chinese individuals and the gender balance across cohorts was found to be comparable (SG90: 59% female, T2D: 52% female, SPMP: 60% female, CPE: 60% female). Age distributions for all cohorts can be found in **Supplementary Figure 14**.

### DNA library construction and sequencing

PowerSoil DNA Isolation Kit (MO Bio Laboratories) was used for the extraction of DNA from stool samples. Minor modifications to the manufacturer’s protocol were made (double the volume of C2, C3 and C4 buffers was added, and the duration of the centrifugation step was extended to twice the original duration). Purified DNA was eluted in 80 µL of C6 solution. DNA libraries were prepared using 50 ng of extracted DNA re-suspended in a volume of 50 µl. This was subjected to shearing using Adaptive Focused Acoustics^TM^ (Covaris) with the following parameters – Duty Factor: 30%, Peak Incident Power (PIP): 450, 200 cycles per burst, Treatment Time: 240s. Sheared DNA was cleaned up with 1.5× Agencourt AMPure XP beads (A63882, Beckman Coulter) followed by end-repair, A-addition and adapter ligation using the Gene Read DNA Library I Core Kit (Qiagen) according to the manufacturer’s protocol. Custom barcode adapters were used instead of GeneRead Adapter I Set for adapter ligation (**Supplementary Table 3**). Before enrichment, DNA libraries were cleaned twice using 1.5× Agencourt AMPure XP beads (A63882, Beckman Coulter) using the protocol from Multiplexing Sample Preparation Oligonucleotide kit (Illumina). Enrichment PCR was carried out with PE 1.0 and custom index-primers for 12 cycles. DNA Libraries were prepared with Agilent DNA1000 Kit (Agilent Technologies) by pooling equimolar concentrations and quantified using Agilent Bioanalyzer. A pilot set of 50 DNA libraries were sequenced on an Illumina HiSeq 2500 and the rest were sequenced on an Illumina HiSeq X instrument (**Supplementary Data File 1**) generating >17 million 2×101 bp reads on average per library. The sequencing protocol and the median sequencing depth of other cohorts can be found in **Supplementary Table 4.**

### Read pre-processing and profiling

Illumina shotgun metagenomic sequencing reads were processed using a Nextflow pipeline (https://github.com/CSB5/shotgunmetagenomics-nf). Briefly, raw reads were filtered to remove low quality bases and adapter sequences were removed using fastp^115^ (v0.20.0) with default parameters. Human reads were removed by mapping to the hg19 reference using BWA-MEM^116^ (v0.7.17-r1188, default parameters) and samtools^117^ (v1.7). The remaining reads were used for taxonomic profiling using MetaPhlAn2^118^ (v2.7.7, default parameters). Functional profiles for the metagenomes were obtained using HUMAnN2^119^ (v2.8.1). For all statistical tests, Benjamini-Hochberg’s false discovery rate method was used to correct for multiple testing at a significance threshold of 5%.

### Statistical analysis with taxonomic profiles

Taxonomic profiles were corrected for batch effects with MMUPHin^120^ using age group as a covariate. MMUPHin is a batch correction method that employs an empirical Bayes approach to model read counts with respect to batch variables and biologically relevant covariates (age group in our case). It then gives as output batch-corrected count data which aims to retain the effects of biologically relevant covariates. For batch correction, we considered samples from the SG90 and CPE cohorts as belonging to the same batch due to their similarity in DNA extraction and library preparation methods i.e. the major sources of variation in metagenomic profiles^121^ (**Supplementary Table 4**). To assess if differences in sequencing platform and depth could affect this assumption, we performed differential abundance analyses using MaAsLin2^122^ (v 1.10.0) (correcting for age) between (i) subsets of the SG90 cohort sequenced on the two different platforms (Illumina HiSeq 2500 and Illumina HiSeq X), and (ii) down-sampled version of the CPE dataset with median sequencing depth matching the SG90 cohort compared to the original CPE dataset. In both analyses we did not identify any differentially abundant taxa between the respective groups, supporting our treatment of the two cohorts as the same batch. We conducted a sanity-check on batch corrected profiles to confirm that cohort effects were reduced (**Supplementary Figure 15**). Species level relative abundances (>0.1%) were then used to compute alpha diversity indices (Shannon and Simpson), Pielou’s evenness and the beta diversity index (Bray-Curtis distance) using the R package vegan^123^ and visualized using ggplot2^124^. Robustness of computed metrics to variations in sequencing depth was confirmed based on sub-sampling and correlation analysis (**Supplementary Figure 16**). For sub-sampling analysis, we selected a subset of 30 samples each with at least 15 million reads, and randomly subsampled 7, 14 and 21 million reads from each of these samples. We then generated the taxonomic and pathway profiles for each of these sequencing depths using the same Nextflow pipeline and computed Spearman correlation values for alpha diversity metrics and pathway counts with the values obtained originally (**Supplementary Figure 16**). Ordinal logistic regression in R was used to test for statistical significance in relation to different age groups. A generalized linear model (GLM) approach was used to test associations between the relative abundance for each species (SA) and age as a continuous variable accounting for covariates.

Demographic and clinical covariates for the analysis were first screened based on previously reported studies^68, 80, 125^ and then through Spearman correlation. We then selected and included only variables that were not highly correlated with one another, retaining only one of the covariates if their correlation was greater than 0.9. The covariates used in this analysis included gender (G), body mass index (BMI), fasting blood glucose (FBG), triglycerides (TGL), total cholesterol (TC) and high-density lipoprotein (HDL) levels (i.e. with the formula: Age ∼ SA + G+ BMI + FBG + TGL + TC + HDL). Only taxa with >50 non-zero values were tested to avoid spurious associations. Missing clinical covariates were imputed through an Expectation Maximization algorithm^126^.

To assess reproducibility of taxonomic associations, three distinct approaches were used: (i) GLM analysis using all four cohorts with batch correction as described above, (ii) GLM analysis using the original data for the SG90 and CPE cohorts as they were similarly processed, and (iii) Trend analysis using all four cohorts with the Cochran-Armitage test^127^ after conversion of relative abundance data into presence-absence values (cutoff of 0.1%). Microbial associations in same direction found in two out of the three methods were considered reproducible with the reported p-values being the median across all methods. In order to further test the robustness of associations, parameters and options were varied for all three approaches including normalization technique (Total Sum Scaling -TSS and Cumulative Sum Scaling -CSS), association analysis technique (ANCOM-BC^128^, MaAsLin2^122^) and relative abundance cutoffs (0.05%-1%; **Supplementary Figure 3**). To obtain a “high stringency” list of microbial associations we restricted to taxa within the SG90 + CPE cohort that are either (i) reported by all four conditions or (ii) reported by at least three conditions, in which case the presence-absence tests are required to report an association at all tested cutoff values (**Supplementary Figure 3**). This list avoids the possibility of a false positive that emerges solely due to batch correction artefacts. The final taxa in this list were visualized using ggplot2 in R. CCREPE^129^ was used to compute Spearman correlation values and identify bacterial species with strong co-occurrence patterns in different age groups (ρ>0.2, p-value<0.05).

To further assess the robustness and reproducibility of our associations, we identified microbial associations across six additional cohorts of Asian and Western origin (**Supplementary Figure 5**). Metagenomic taxonomic profiles from stool samples for four of these cohorts (labelled as YachidaS_2019^39^, AsnicarF_2017^40^, BritoIL_2016^41^ and LLD_2016^42^) were obtained from curatedMetagenomicData^114^. The subjects in these cohorts were healthy and were selected for not having antibiotic usage. Microbial associations and taxonomic profiles for two other cohorts (labelled as XuWuZhu_2022^43^ and ElderMet_2020^17^) were obtained from their respective publications. We then employed a similar GLM framework (with the formula: Age ∼ SA + G) as outlined previously to identify microbial associations with age. Microbial associations from our results are considered ‘replicated’ or ‘validated’ against other cohorts based on statistical significances (FDR-adjusted p-value<0.05 or p-value<0.05 respectively).

### Pathway and gene-level analysis

The HMP Unified Metabolic Analysis Network (HUMAnN2)^130^ pipeline was used to determine the relative abundance of microbial pathways in different gut metagenomes. The default Kyoto encyclopedia of genes and genomes (KEGG)^131^ catalog was used as the pathway reference. Unstratified relative abundance values for SG90, CPE and SPMP were integrated with HUMAnN2 results for T2D from curatedMetagenomicData^114^ for shared pathways, and significant differentially abundant pathways across age groups were determined based on linear discriminant analysis with LEfSe^132^ (p-value<0.05 and LDA score >3).

For gene-level analysis of metabolic pathways involved in butyrate production, EC numbers of enzymes were mapped to corresponding UniRef90^133^ identifiers from the HUMAnN2 output. The list of butyrate producing species, genes, and enzymes (EC numbers) in butyrate production pathways were obtained from previously published studies^45, 46^. Using this list, the butyrate production genes of interest from our dataset were obtained using a three step process: (i) UniRef90 identifiers obtained from HUMAnN2, were first cross-referenced with the KEGG database to obtain a list of KEGG orthology (KO) groups, (ii) EC numbers obtained previously were matched with the KO groups (**Supplementary Data File 7**), and (iii) the KO groups were cross-referenced with UniRef90 to obtain the specific butyrate producing genes of interest. For every gene, read counts were obtained by taking the sum for all corresponding UniRef90 identifier-associated values and statistical analyses were performed using R with visualizations generated using ggplot2^124^.

### Metagenomic assembly

Microbial reads from the SG90 cohort were assembled and binned following an in-house snakemake pipeline. Briefly, the reads were assembled using MEGAHIT^134^ (v1.2.9). Contigs were binned using MetaBAT2^135^ (v2.15) based on contig coverage information generated with minimap2^136^ (v2.26). Assemblies were evaluated based on MIMAG^137^ definitions, with contamination, completeness, and N50 values obtained from CheckM2^138^ (v1.0.2), and non-coding RNA identified using barrnap (https://github.com/tseemann/barrnap) (v0.9) and tRNAScan-SE^139^ (v2.0.12). Potential cases of chimerism were detected using GUNC^140^ (v1.0.5). All software were used with default arguments. Low-quality MAGs were removed from downstream analysis and taxonomic classification of MAGs at the species level was obtained using GTDB-Tk^141^ (v2.3.2) ‘classify_wf’ method (GTDB version r214). Genomes were annotated using Prokka^142^ (v1.14.6) and the EC numbers associated with each gene were extracted. Metabolic pathways were obtained using MinPath^143^ (v1.6). Reachability analysis was carried out using MetQuest^144^ (v0.1.33) using the bipartite graphs constructed from these metabolic pathways. The initial seed list for the reachability analysis included source metabolites L-aspartate for L-lysine biosynthesis III and L-lysine for L-lysine to butyrate pathway.

### Mouse experiments

All the mouse experiments were carried out with the approval of the Institutional Animal Care and Use Committee (IACUC) of National University of Singapore (NUS) under the R21-0135 protocol.

Two independent groups of 18 month old C57BL/6 mice (n=20 each, regular chow diet) were housed separately (maximum of four mice per cage). Mice in the control group were on the regular chow diet while those in the healthy aging group were switched to a diet containing 4% calcium alpha-ketoglutarate, with stool samples collected at baseline and 3 months after diet switch for metagenomic analysis. Genomic DNA was extracted from mouse stools using QIAamp PowerFecal Pro DNA Kit (Qiagen), according to manufacturer’s instructions. DNA was quantified on a Qubit Fluorometer using the Qubit dsDNA BR Assay Kit (ThermoFisher Scientific). Purified genomic DNA (50 ng) was used for library construction steps using NEBNext® Ultra™ II FS DNA Library Prep Kit according to manufacturer’s instructions. Finally, each library sample was assessed for quality based on fragment size and concentration using the Agilent D1000 ScreenTape system, with samples adjusted to identical concentrations by means of dilution and volume-adjusted pooling. The multiplexed sample pool was paired-end (2×151 bp) sequenced on an Illumina HiSeq X Ten system.

Mouse gut metagenomes were annotated using eggNOG-mapper^145^ (v2.1.9). Briefly, reads were mapped against eggNOG protein database (eggnog 5.0^146^) using DIAMOND^147^ in blastx mode. The default KEGG^131^ catalog was used as the pathway reference. For gene-level analysis of metabolic pathways involved in butyrate production, the corresponding KO IDs of each gene were used from the eggNOG-mapper’s output. For every gene, read counts were obtained by taking the sum for all corresponding KEGG values. Gene normalized abundances were measured as logarithm of counts per millions [log(CPM + 1)] to adjust for differences in sequencing depths. Pathway normalized abundances were calculated by taking the sum of all of gene counts for each pathway and log-transformed and normalized to sequencing depth as log(CPM + 1). Statistical tests for groups comparisons were done using Wilcoxon rank-sum test (FDR adjusted p-value < 0.05 considered to be significant) and were performed using R with visualizations created using ggplot2.

### Metabolic network analysis

Metabolic support index (MSI) values for the species identified in **Table 2** in relation to other gut microbial species were obtained based on metabolic network analysis, as described previously^148^. Briefly, MSI uses network flow analysis to quantify the extent of microbial metabolism that is enabled by the presence of other species in the community. The metabolic support network was visualized as a directed graph using Cytoscape^149^ (v3.8.0). Species that are known butyrate producers^45, 46^ and receive the most metabolic support (based on in-degree of nodes) were highlighted in the network. The combined relative abundances of organisms supporting these butyrate producers were compared across age groups using the Wilcoxon test.

### Phenotypic association analysis

Associations of microbial species with clinical phenotypic markers were identified based on linear regression using ‘glm’ function in R, with age and other markers serving as covariates for adjustment. Only species present in at least ten samples were considered for this analysis. Phenotypic markers available in two datasets (T2D and SG90) included body mass index (BMI), fasting blood glucose, triglycerides, total cholesterol, high-density lipoprotein (HDL) and low-density lipoprotein (LDL) levels. For hs-CRP, AST, and ALT levels, gait speed, right handgrip strength, left handgrip strength, sleep duration, MMSE and free thyroxine levels, the associations were only tested for SG90 samples (without batch-correction) where this data was available.

## Data and source code availability

Shotgun metagenomic sequencing data are available from the European Nucleotide Archive (ENA – https://www.ebi.ac.uk/ena/browser/home) under project accession number PRJEB49124. Source code for scripts used to analyze the data are available in a GitHub project at https://github.com/CSB5/SG90.

---

**Supplementary Figure 1:**
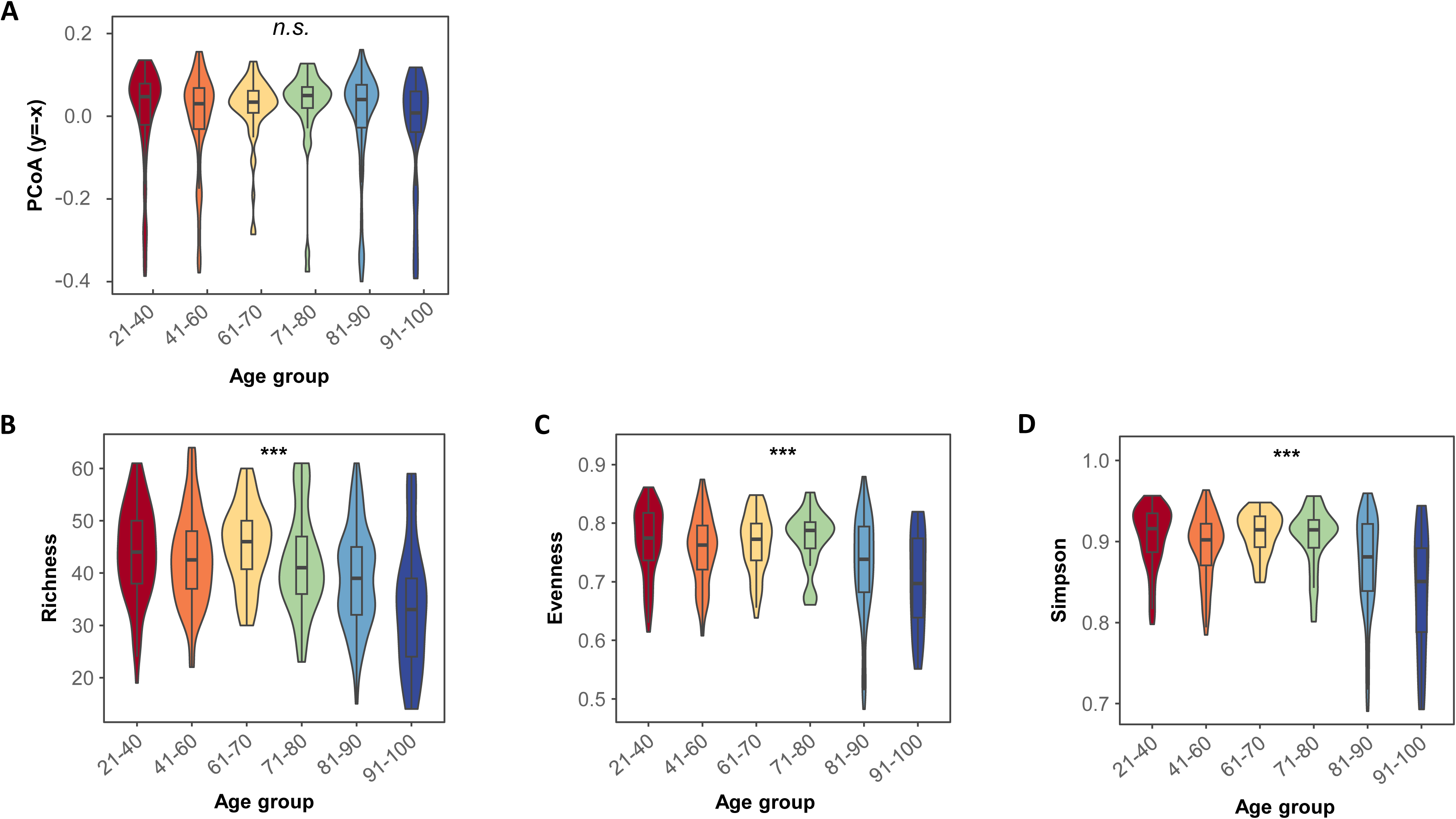
Variation in gut microbiome beta and alpha diversity metrics across age groups. (A) Violin plot showing the variation of beta diversity on the y=-x axes corresponding to the PCoA plot in Figure 1A. Violin plots showing species-level (A) Richness, (B) Evenness and (C) Simpson diversity index, across age groups. All three indices were found to be significantly different between age groups (ordinal logistic test; “***”: p-value<0.001). Data points that are either less than Q1 – 1.5 × IQR or more than Q3 + 1.5 × IQR, where Q1, Q3 and IQR refers to the first quartile, third quartile and interquartile range, respectively, were removed for this analysis to reduce the impact of outliers.

**Supplementary Figure 2:**
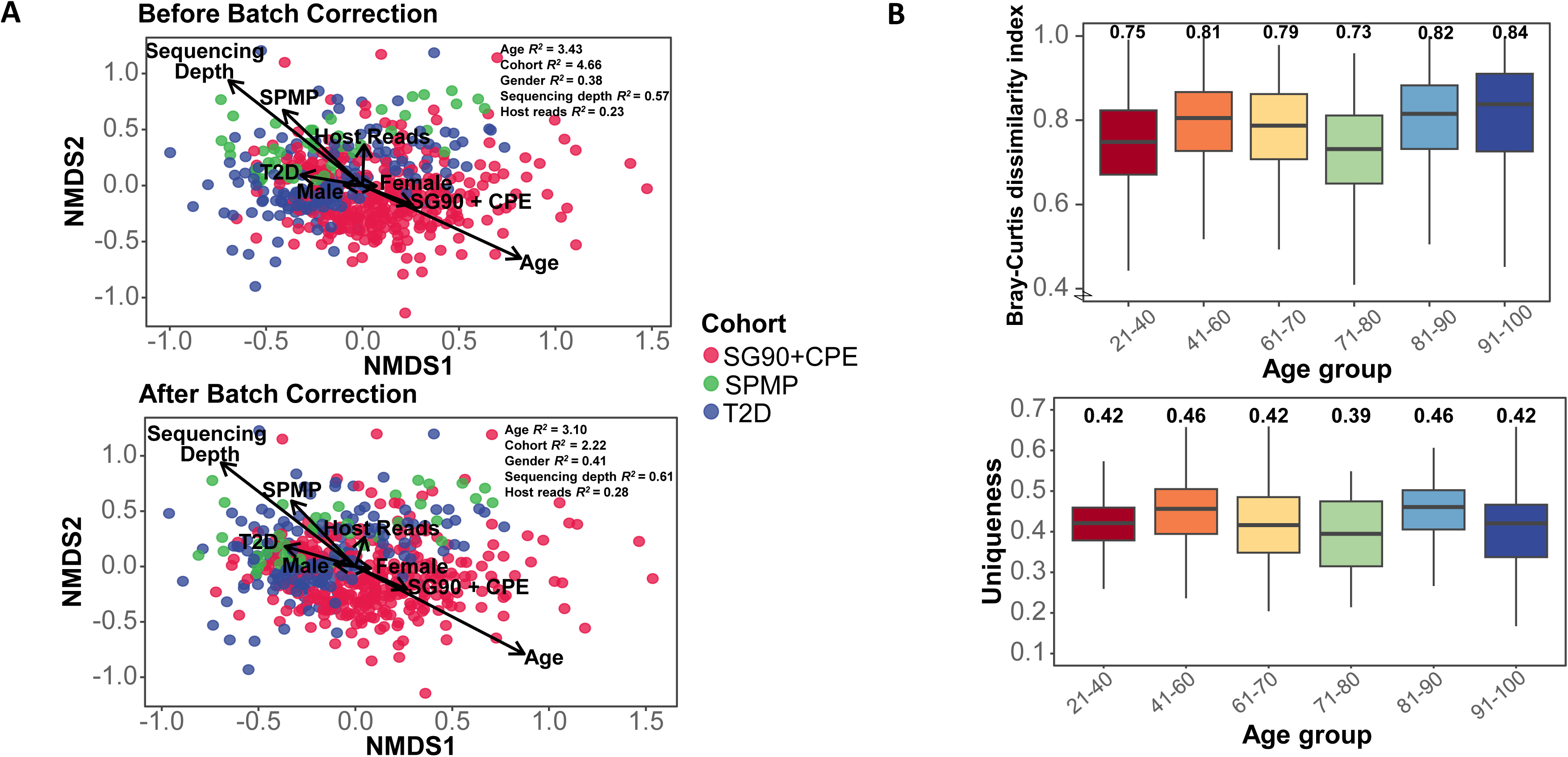
Variation of beta diversity metrics. A) Nonmetric Multidimensional Scaling (NMDS) plots showing the distribution of samples for each cohort before (top) and after (bottom) batch correction. Vectors for each of the confounding variables were obtained through envfit analysis demonstrating the relationship between the ordination axes and the variables. Length of the vector indicates strength of the relationship, and the direction points to the steepest increase corresponding to the variable. Inset values for R^2^ are from PERMANOVA analysis using Bray-Curtis dissimilarity. B) Box plots showing the species-level Bray-Curtis dissimilarity index (top), Uniqueness (bottom) across age groups. Uniqueness was defined as the minimum value of Bray-Curtis dissimilarity index for each individual in all boxplots, the center line represents the median, box limits represent upper and lower quartiles, and whiskers represent minimum and maximum values. Median values are shown on top of the plot.

**Supplementary Figure 3:**
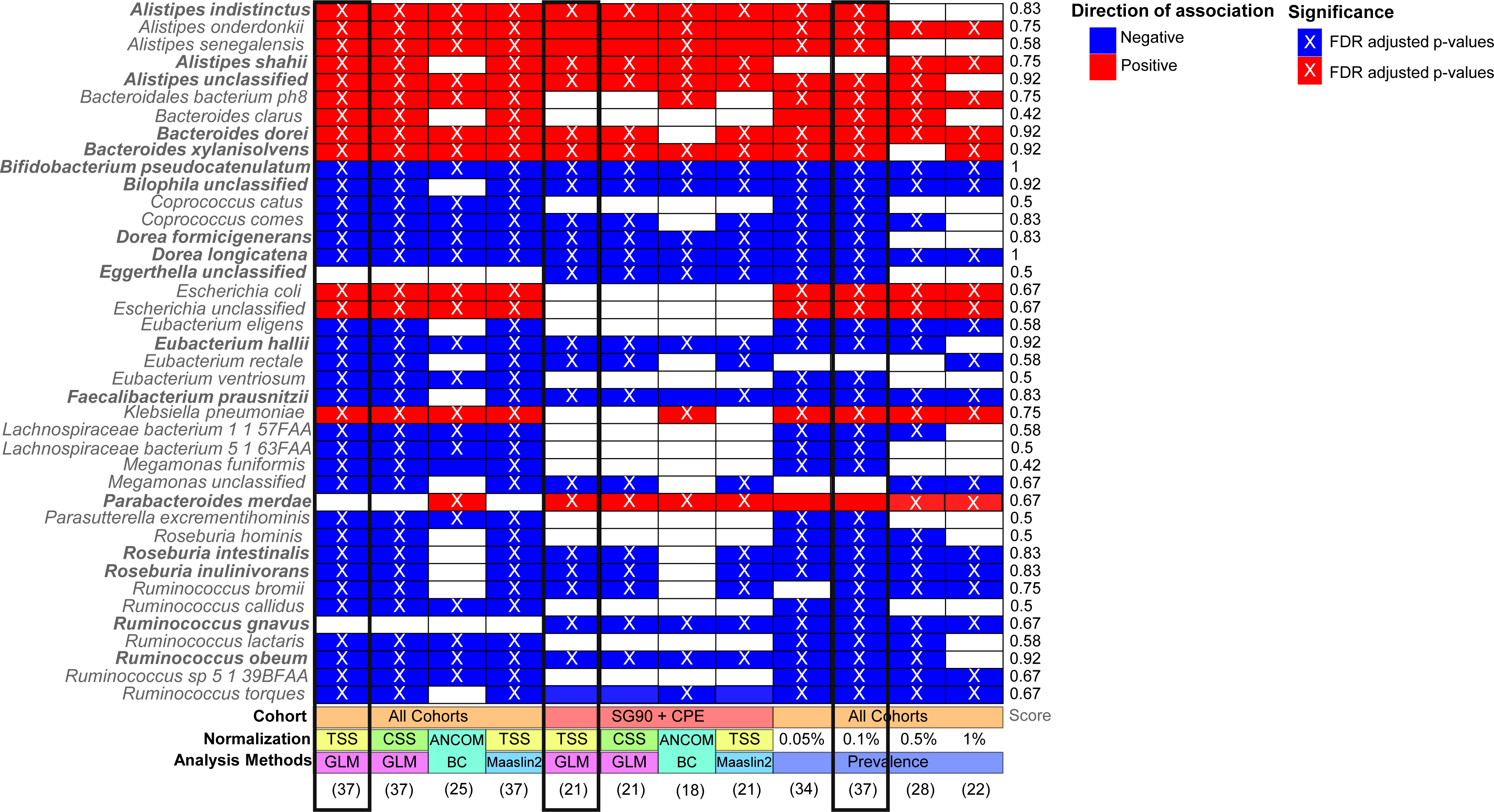
Assessing the reproducibility and robustness of microbial associations with age. Heatmap showing microbial species associations with age, where (i) reproducibility was assessed through 3 distinct primary analysis approaches (black boxes) and (ii) robustness was assessed by varying their parameters, normalization and analysis settings (all columns; see **Methods)**. The robustness score (last column) captures the frequency with which a taxa was observed to be significantly associated with age across all tested methods and settings (FDR adjusted p-value<0.05). Note that all but two associations had robustness score greater than or equal to 50%. Numbers in parentheses below each column indicate the number of significant associations for each method. Red and blue colored cells represent positive and negative associations, respectively. Taxa found in the high-stringency list are in bold face. Colored cells marked with ‘X’ correspond to FDR adjusted p-value<0.05, while cells without an ‘X’ correspond to unadjusted p-value<0.05.

**Supplementary Figure 4:**
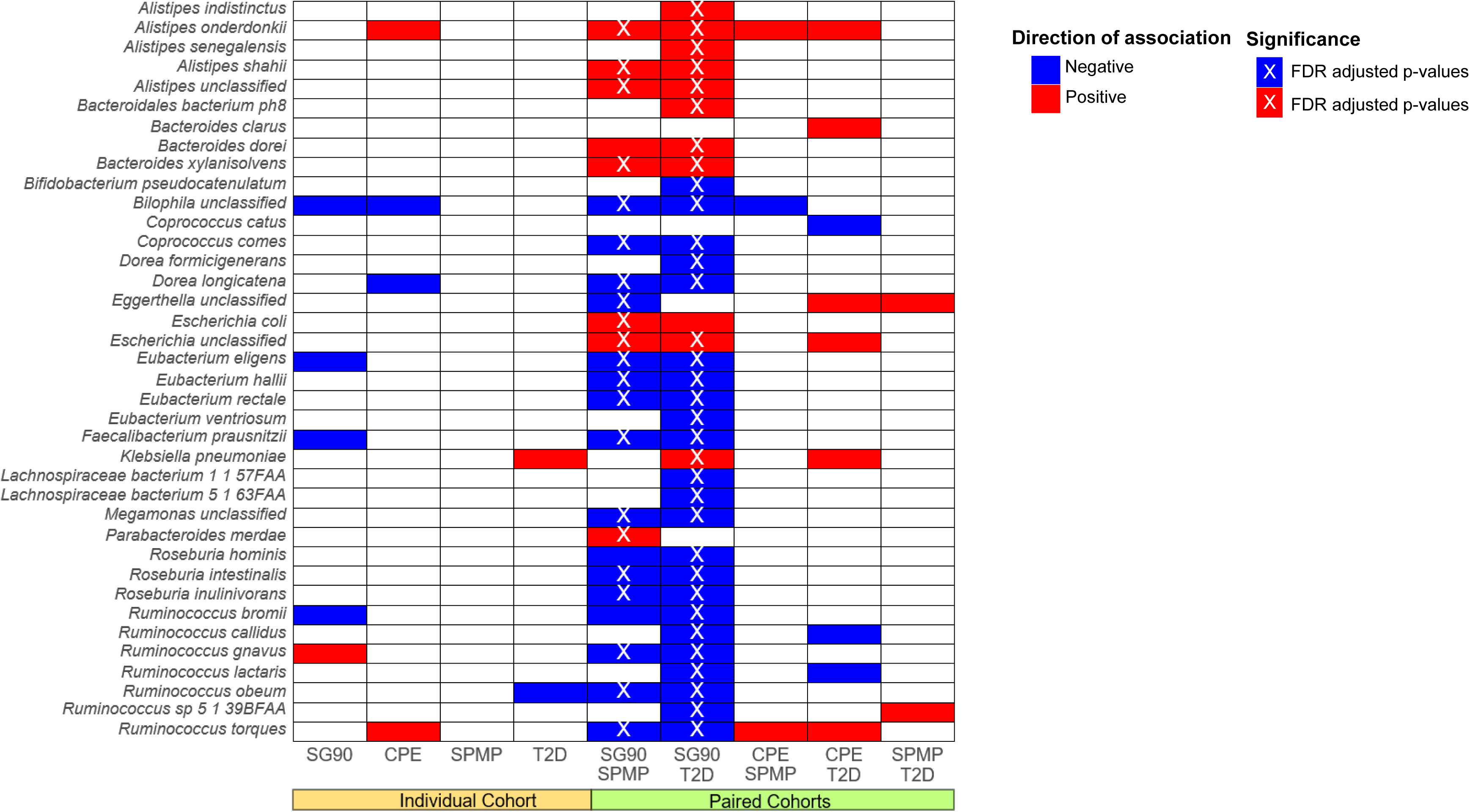
Associations within individual and paired cohorts. Heatmap showing microbial species associations with age identified using independent and paired cohort analysis. Red and blue colored cells represent positive and negative associations, respectively. Colored cells marked with ‘X’ correspond to FDR adjusted p-value<0.05, while cells without an ‘X’ correspond to unadjusted p-value<0.05. As can be seen here, while some associations are identified in individual cohorts (n=10), joint analysis of SG90 with a cohort of younger individuals is essential to identify many more associations (n=38). These association are unlikely to be purely a function of batch effects as very few associations are detected when other pairs of cohorts are analyzed together (e.g. n=3, using CPE and SPMP), where these associations are also not significant after FDR adjustment.

**Supplementary Figure 5:**
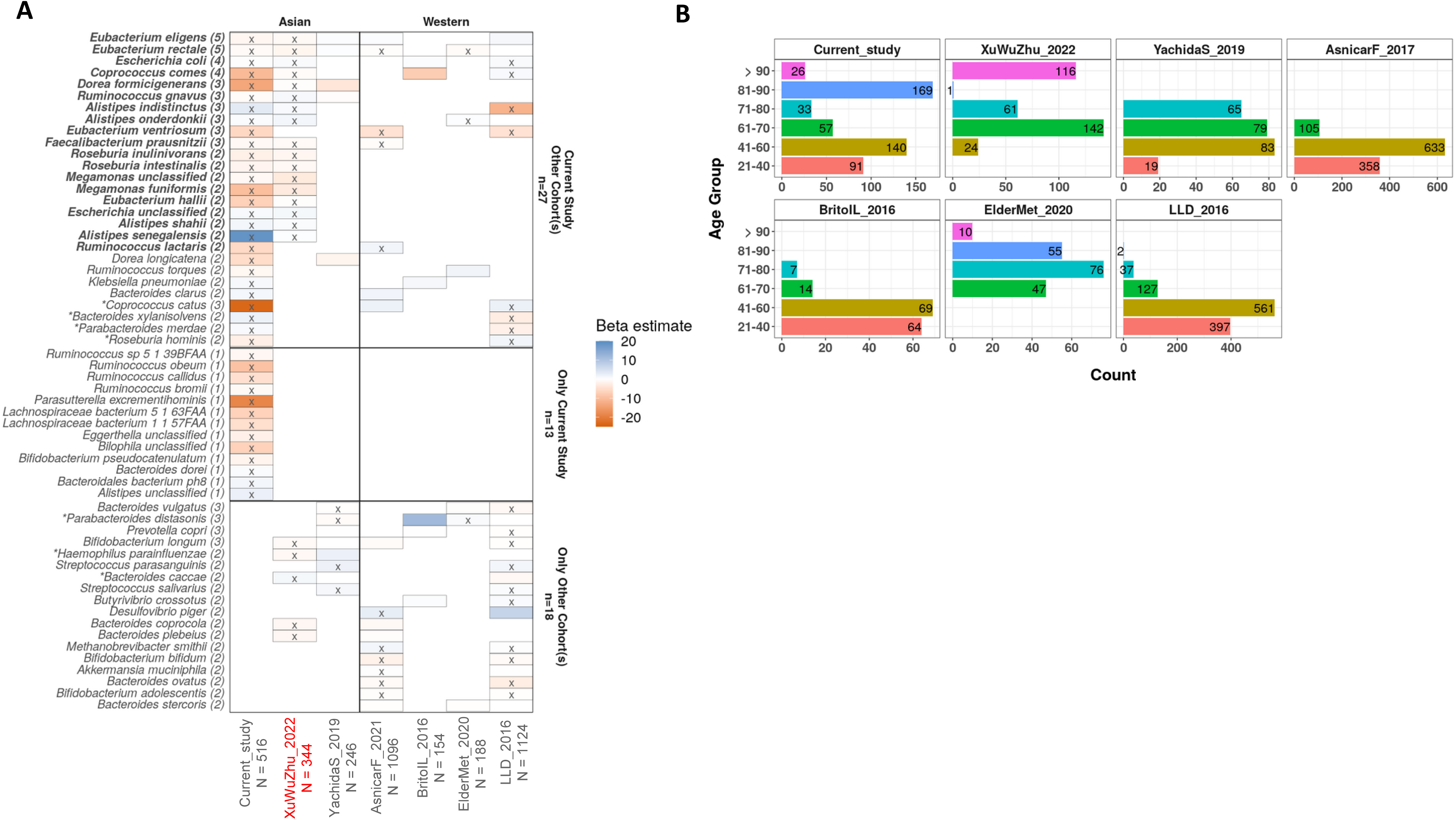
Associations within other cohorts. A) Heatmap showing taxonomic associations with age across Asian and western cohorts. Colored cells marked with ‘X’ correspond to FDR adjusted p-value<0.05, while cells without an ‘X’ correspond to unadjusted p-value<0.05. Colors of the boxes represent whether the organism is positively (blue) or negatively (red) associated with age based on beta estimate. The numbers in the bracket represent the number of times a particular taxa is validated in other cohorts (p-value <0.05). Taxa in bold face indicate those that are independently replicated in other cohorts (FDR adjusted p-value <0.05). Taxa marked with ‘*’ are those whose the directionality of association is different across cohorts. The cohort ‘XuWuZhu_2022’ is colored red as the association results were obtained from the original manuscript and are based on a Wilcoxon rank-sum test instead of GLM analysis. B) Bar plot showing the number of subjects (indicated in every bar) for different cohorts in each age group. The number of subjects in each age group is calculated by including the lower and upper end of the range.

**Supplementary Table 1:**
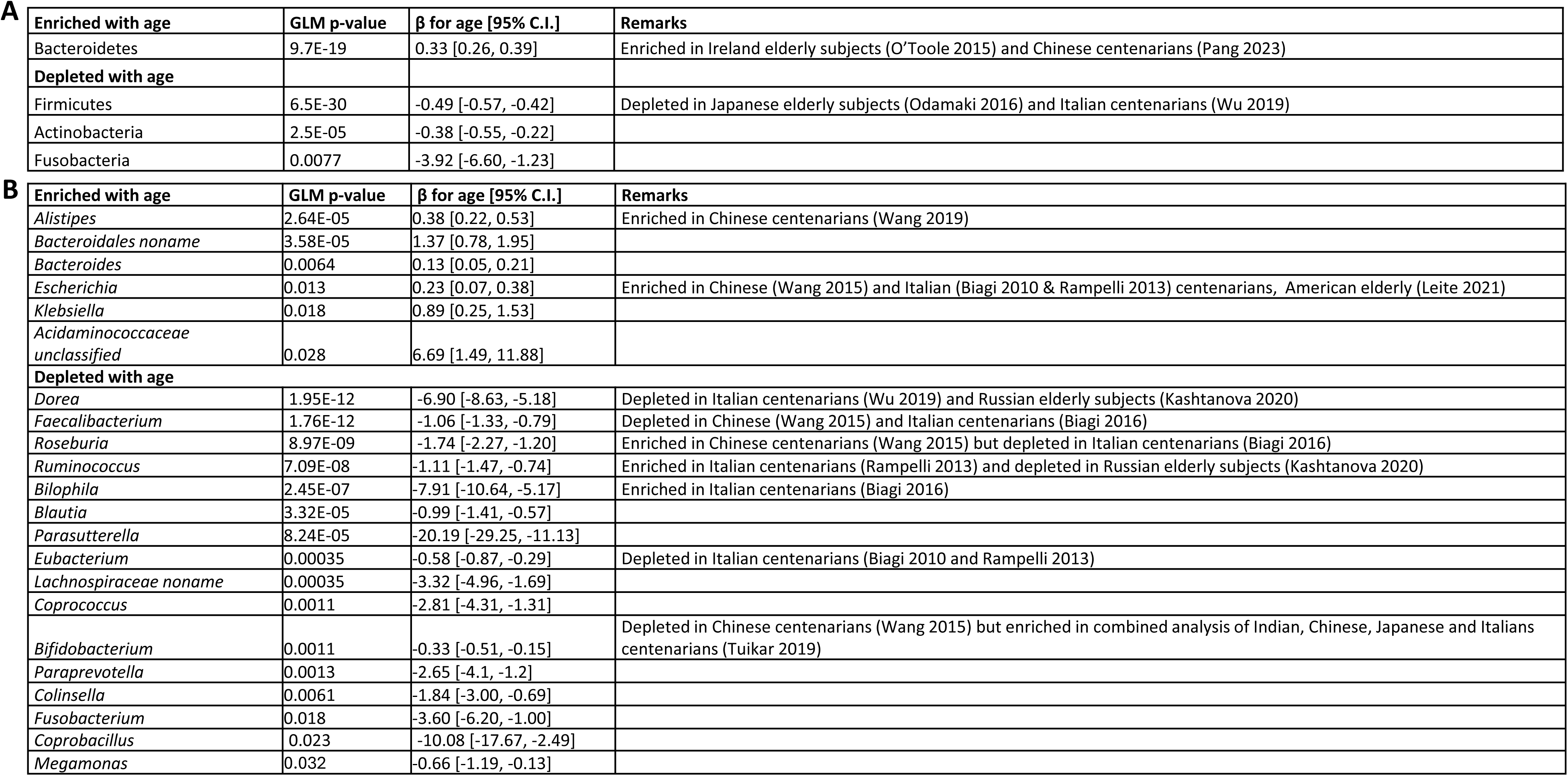
Higher-level taxonomic associations with age. At the (A) phylum and (B) genus level, the table reports taxa that are statistically significant with FDR-adjusted p-values≤0.05 from GLM test with co-variate adjustment (see **Methods**). The p-values reported here are the FDR-adjusted p-values from the GLM test. The β coefficient estimates strength of association as change in age (in years) for every 1% increment of taxa relative abundance.

**Supplementary Figure 6:**
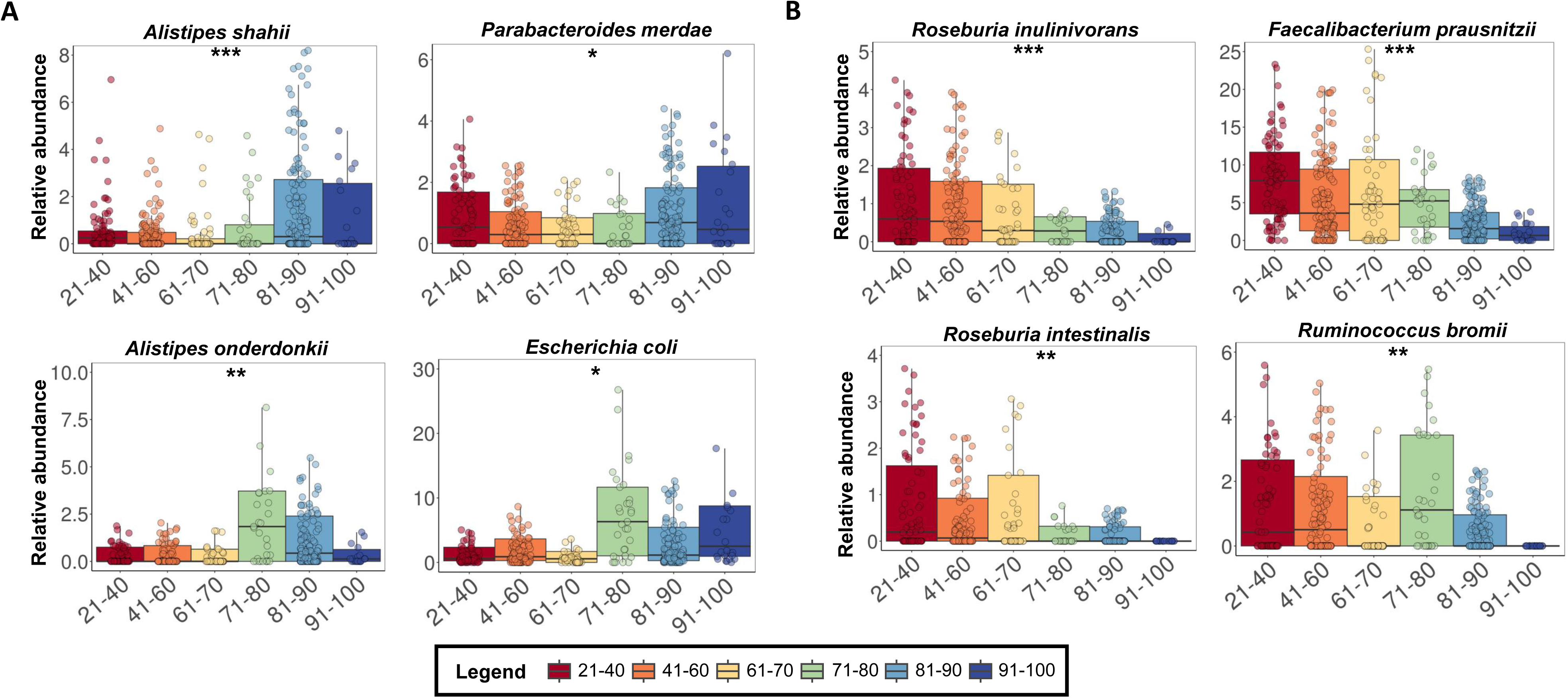
Relative abundance across age groups for key gut microbial species associated with aging. Boxplots showing relative abundances across age groups for (A) species whose abundances increase with age and (B) species whose abundances decrease with age. Median FDR-adjusted p-values from the 3 primary analysis approaches are denoted by “*” (p-value<0.05), “**” (p-value<0.01) and “***” (p-value<0.001). In all boxplots, the center line represents the median, box limits represent upper and lower quartiles, and whiskers represent minimum and maximum values (outlier points are not included in the visualization).

**Supplementary Figure 7:**
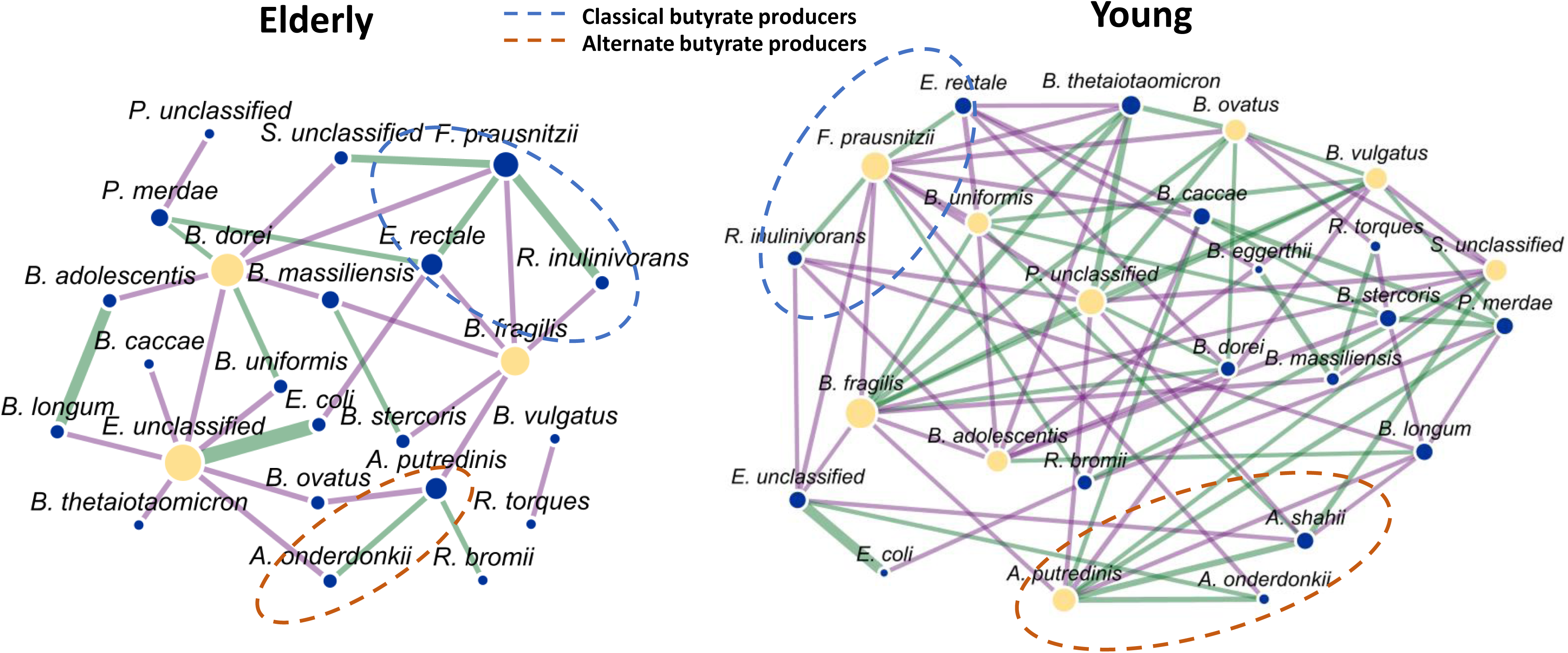
Microbial co-occurrence patterns in elderly and young Asian subjects. Nodes in the network represent microbial species and edges represent significant Spearman correlations between relative abundances of species corresponding to adjacent nodes (|r|≥0.1, p-value<0.05). The network on the left (Elderly) was constructed with samples in the age range 70-100, while that on the right (Young) was constructed with samples in the age range 20-69 to ensure that sufficient number of datapoints were available. Green- and purple-colored edges represent positive and negative correlations respectively. Thickness of the edges are directly proportional to the Spearman correlation value |r|. Blue and orange circles identify the classical and alternate butyrate producer species, respectively. Hubs in the network are shown as yellow-colored nodes. There are 23 nodes, 32 edges, 3 hubs in the elderly network, while the young network has 25 nodes, 80 edges and 9 hubs.

**Supplementary Figure 8:**
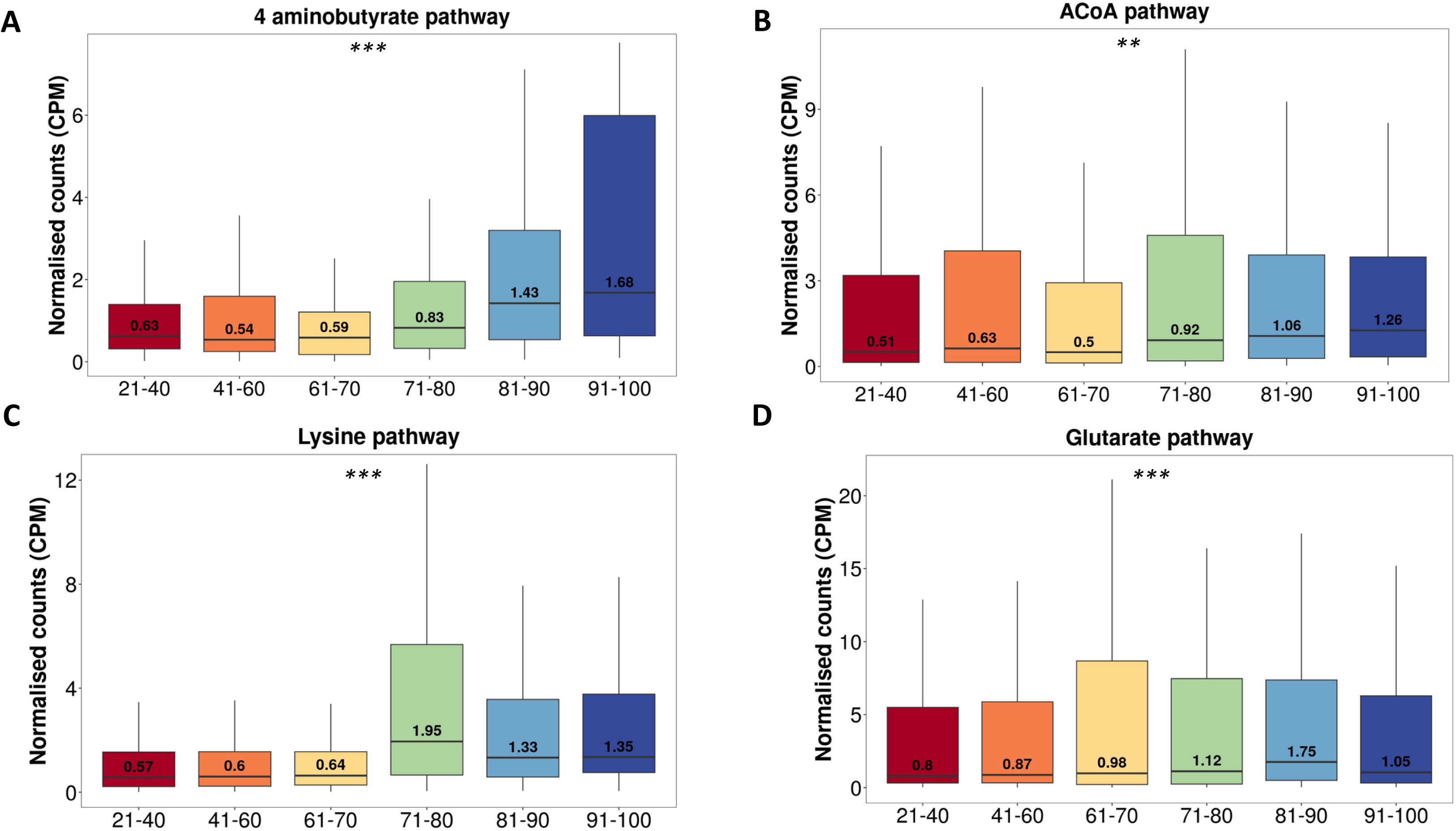
Variation in butyrate production pathways across age groups. Boxplots showing the metagenomic relative abundance of pathways for butyrate production in different age groups. FDR-adjusted p-values from a GLM test for age association are denoted by “**” (p-value≤0.01) and “***” (p-value<0.001). In all boxplots, the center line represents median, box limits represent upper and lower quartiles, and whiskers represent minimum and maximum values (outlier points are not included in the visualization).

**Supplementary Figure 9:**
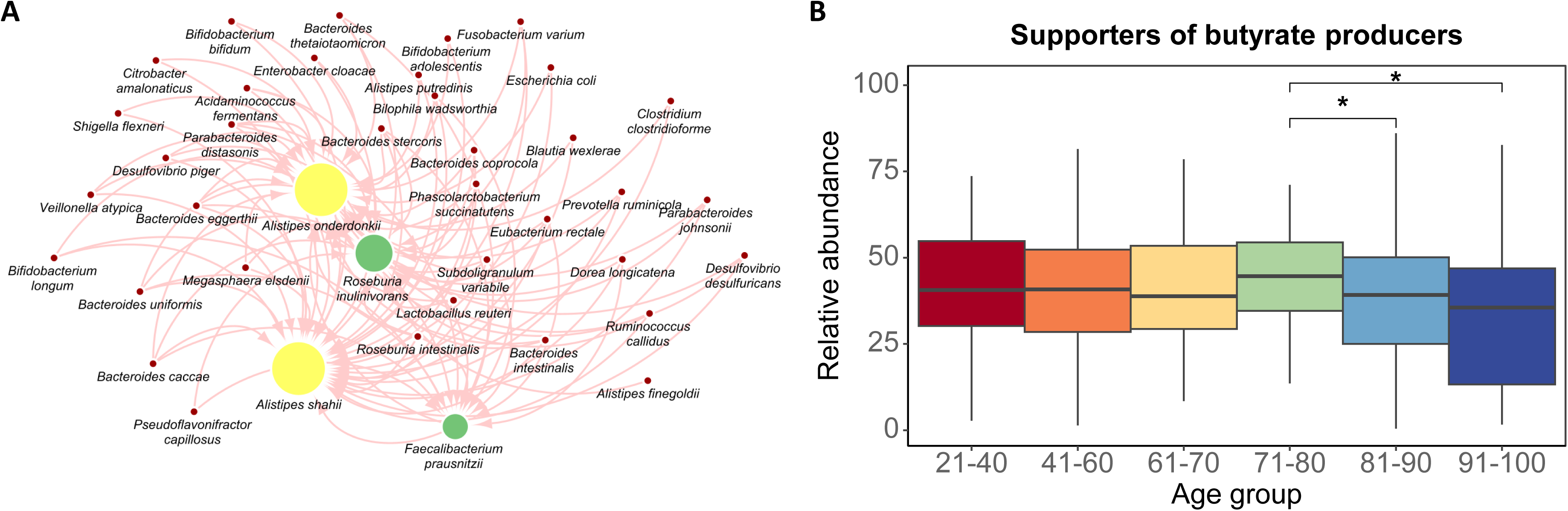
Metabolic support in the gut microbiome for butyrate production. (A) Network figure depicting gut microbial species with high ‘metabolic support index’ for butyrate producers that receive maximum support. Directed edges go from the supporting species to the one that is being supported, and the size of the supported node is proportional to the number of incoming edges. Nodes colored in yellow and green indicate those enriched and depleted with age respectively. (B) Boxplots depicting combined relative abundances of all species that support butyrate producers (from subfigure A) in the gut microbiomes of subjects from various age groups. Wilcoxon test p-values<0.05 are indicated with the star symbol (‘*’). In all boxplots, the center line represents median, box limits represent upper and lower quartiles, and whiskers represent minimum and maximum values.

**Supplementary Figure 10:**
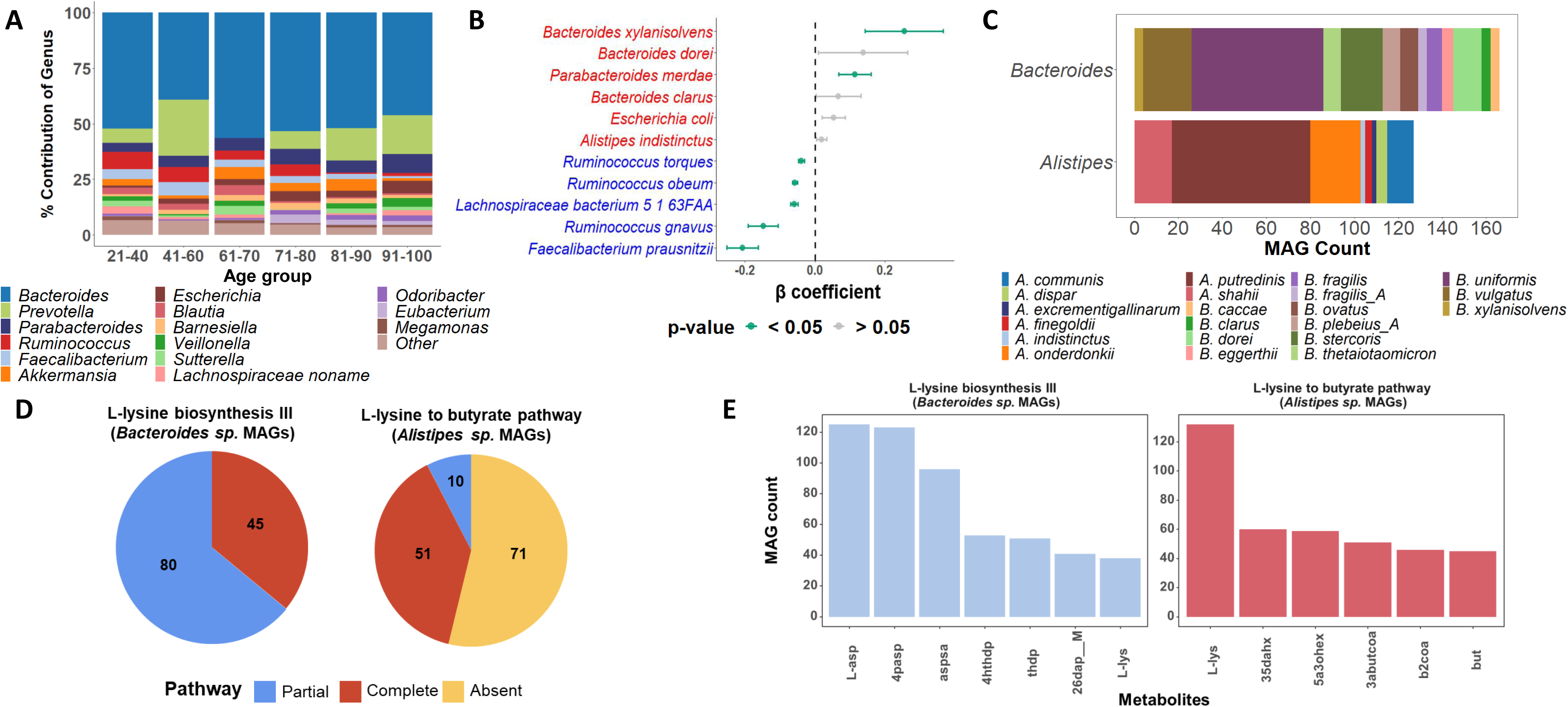
Species contributions to L-lysine biosynthesis III and L-lysine to butyrate pathways. (A) Stacked bar chart showing the relative contributions of genera to L-lysine biosynthesis III pathway. (B) Dot plot showing the variation of beta-coefficient of the age-associated organisms with the abundance of L-lysine biosynthesis III across age. The organisms highlighted in blue are negatively associated with age, while those in red are positively associated. (C) Stacked bar chart showing the diversity of *Alistipes sp.* and *Bacteroides sp.* in the assembled MAGs. (D) Pie chart showing the number of MAGs that were annotated with L-lysine biosynthesis III and L-Lysine to butyrate pathways in *Bacteroides sp.* and *Alistipes sp*. respectively. (E) Bar chart showing the number of times a metabolite is reachable within the MAGs of *Bacteroides sp.* and *Alistipes sp*. in L-lysine biosynthesis III pathway (left) and L-lysine to butyrate pathway (right). X-axis represents the intermediate metabolites in the respective pathways. Metabolites are abbreviated based on BiGG database.

**Supplementary Figure 11:**
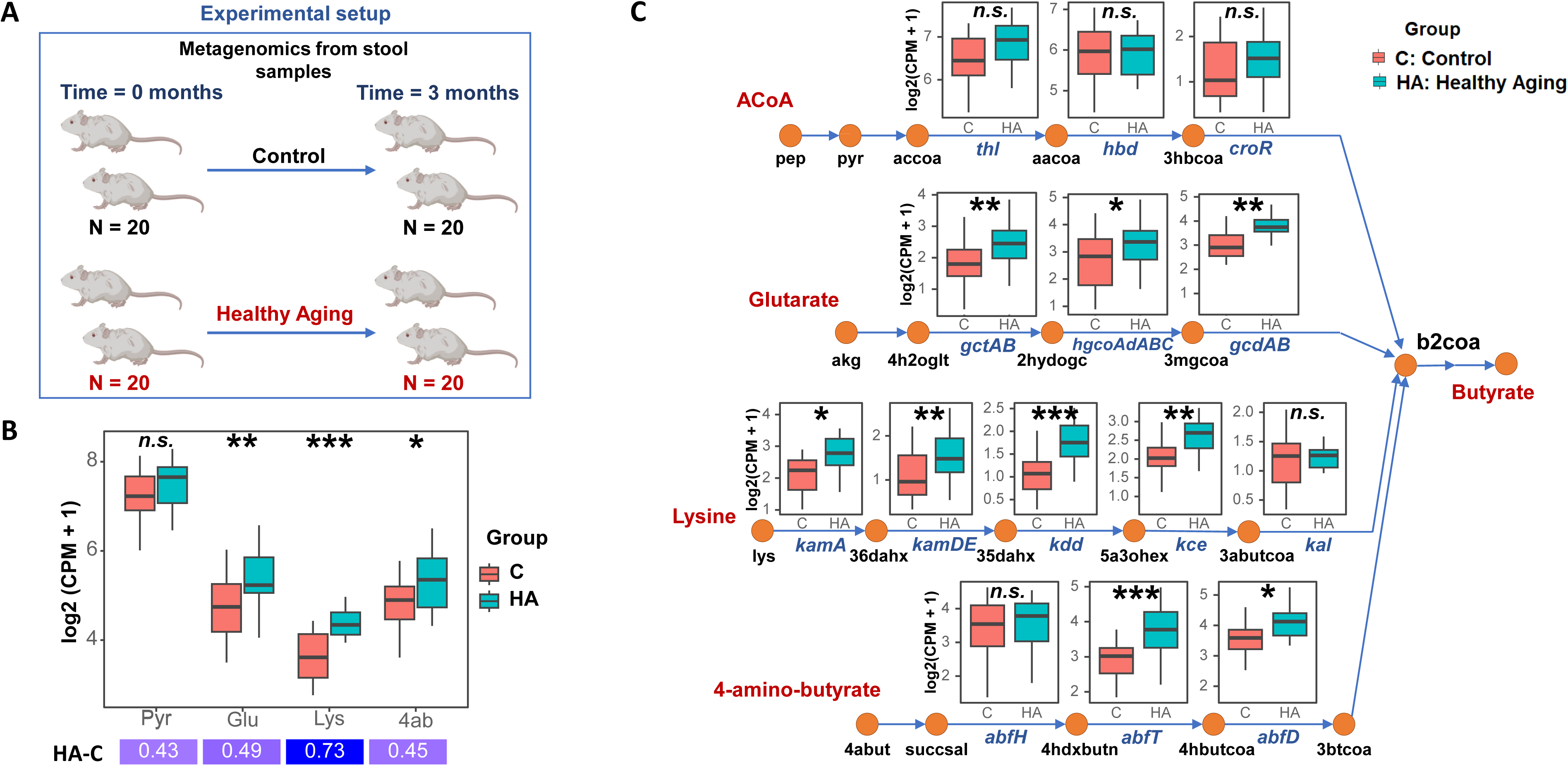
Butyrate production pathways in a healthy aging mouse model. (A) Experimental setup for fecal metagenomic analysis in two groups of aged mice (18 months) with (Healthy Aging -HA) and without (Control -C) CaAKG 4% supplementation. (B) Boxplots showing relative abundance of alternative pathways for butyrate production in the two mouse groups after 3 months of supplementation. C) Boxplots for abundance of various genes in corresponding pathways where nodes represent major metabolites and edge labels represent genes involved in their conversion. In all boxplots, center line represents median, box limits represent upper and lower quartiles, and whiskers represent minimum and maximum values. The labels “*n.s.”,* “*”, “**” and “***” represent FDR-adjusted p-value>0.05, p-value≤0.05, p-value<0.01 and p-value < 0.001.

**Supplementary Table 2:** Reference ranges for various clinical phenotypes. Healthy ranges for various clinical phenotypes that were analyzed in this study.

| Clinical phenotypes | Range |
| --- | --- |
| HDL | 1.0 – 1.5 mmol/L |
| LDL | 2.6 – 3.3 mmol/L |
| Triglyceride | 1.7 – 2.2 mmol/L |
| Total cholesterol | < 5.2 mmol/L |
| Fasting blood glucose | < 6.1 mmol/L |
| High-sensitivity CRP | < 1 mg/L |
| Vitamin B12 | 160 to 950 pmol/L |
| Alanine<br>Aminotransferase | 6 - 66 U/L |
| Aspartate<br>Aminotransferase | 6 - 35 U/L |

**Supplementary Figure 12:**
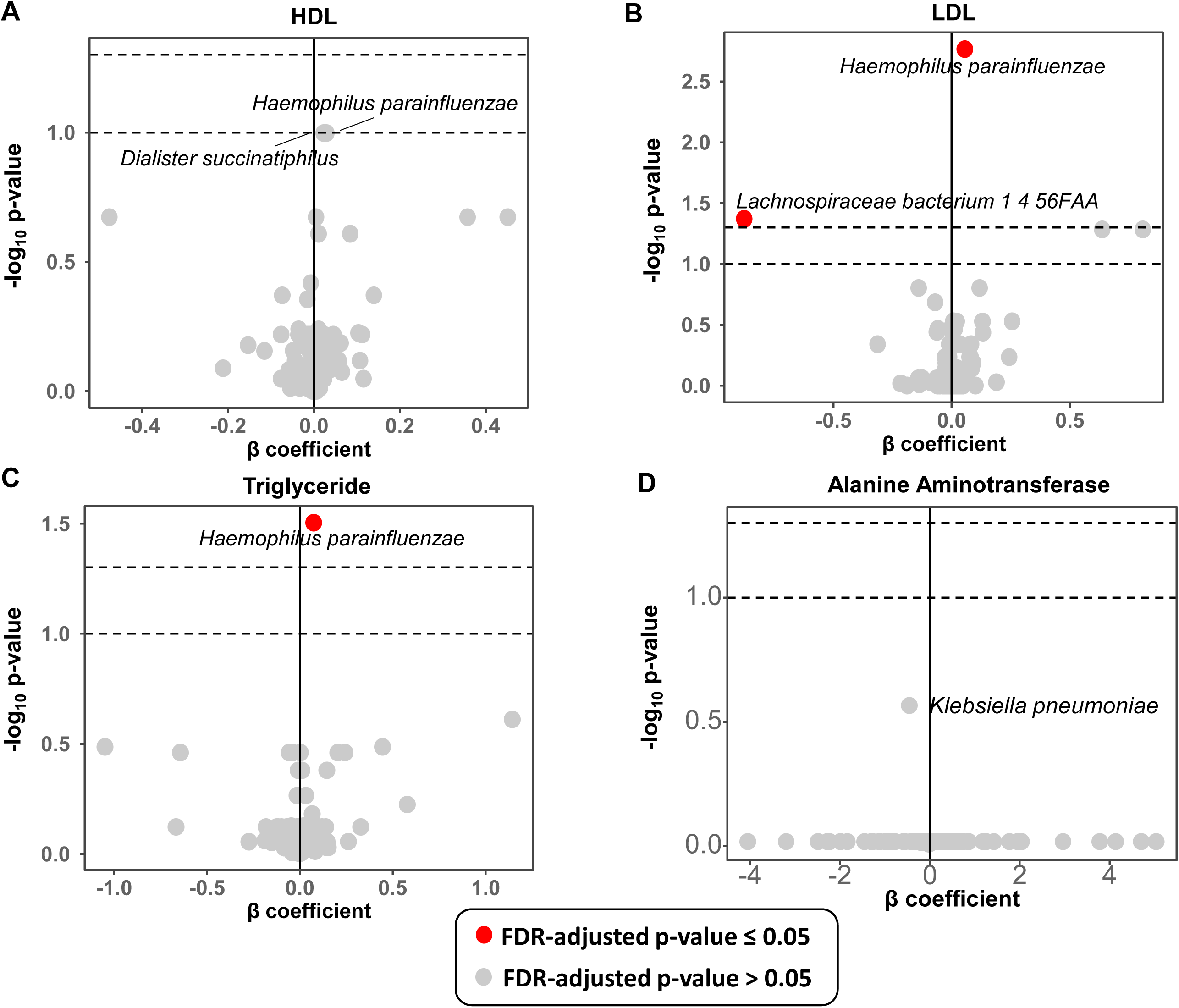
Microbiome associations across various metabolic markers. Volcano plots showing β coefficient on the x-axis and FDR-adjusted –log_10_ p*-*values (GLM test) on the y-axis. Red dots indicate p-value≤0.05 and grey dots indicate otherwise, with dotted lines marking p-value*=*0.05 and p-value*=*0.1 thresholds. (A), (B), and (C) Results for HDL, LDL and Triglyceride levels. (D) Results for Alanine Aminotransferase showing a weak association with *Klebsiella pneumoniae*, compared to the significant association in Figure 4D with AST.

**Supplementary Table 3:**
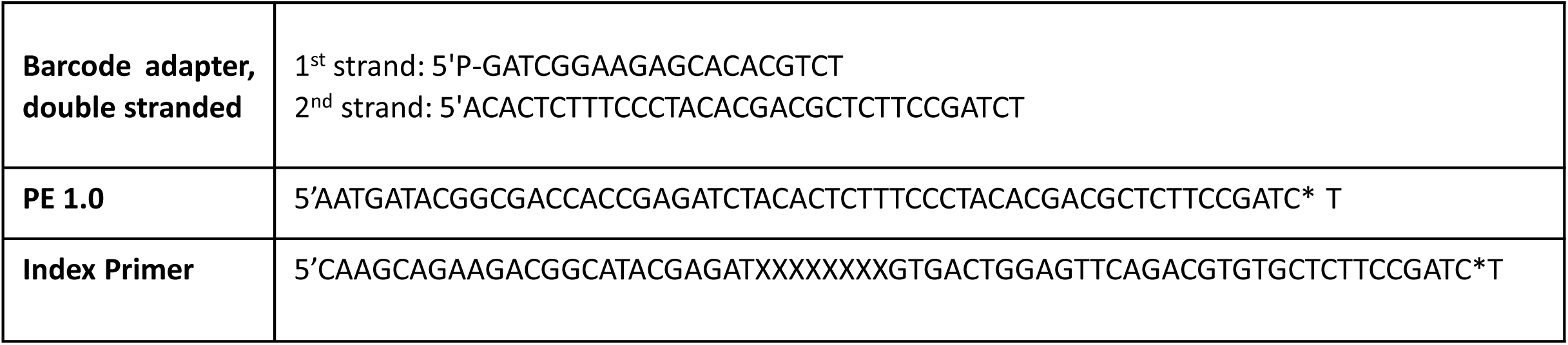
Primers and adapter sequences used in this study.

**Supplementary Table 4:** Experimental methods used in different cohorts for DNA extraction and sequencing.

| Cohort | Sample collection | DNA Extraction kit | Library Preparation kit | Sequencer | Median Sequencing Depth | Median Microbial Reads |
| --- | --- | --- | --- | --- | --- | --- |
| SG90 | Frozen stool samples | PowerSoil DNA Isolation Kit | Gene Read DNA Library I Core Kit | Illumina HiSeq 2500 | 4.3M | 2.9M |
|  |  |  |  | Illumina HiSeq X | 9.1M | 6.4M |
| SPMP | Frozen stool samples | QIAamp Power Fecal Pro DNA kit | NEBNext Ultra DNA Library Prep Kit | Illumina HiSeq4K | 36.7M | 31.2M |
| CPE | Frozen stool samples | PowerSoil DNA Isolation Kit | Gene Read DNA Library I Core Kit | Illumina HiSeq 2500 | 24M | 20.6M |
| T2D | Frozen stool samples | Custom method<sup>1</sup> | Custom method<sup>1</sup> | Illumina GA | 18.2M | NA |

1. Qin, J. *et al.* A metagenome-wide association study of gut microbiota in type 2 diabetes. *Nature* **490**, 55–60 (2012).

**Supplementary Figure 13:**
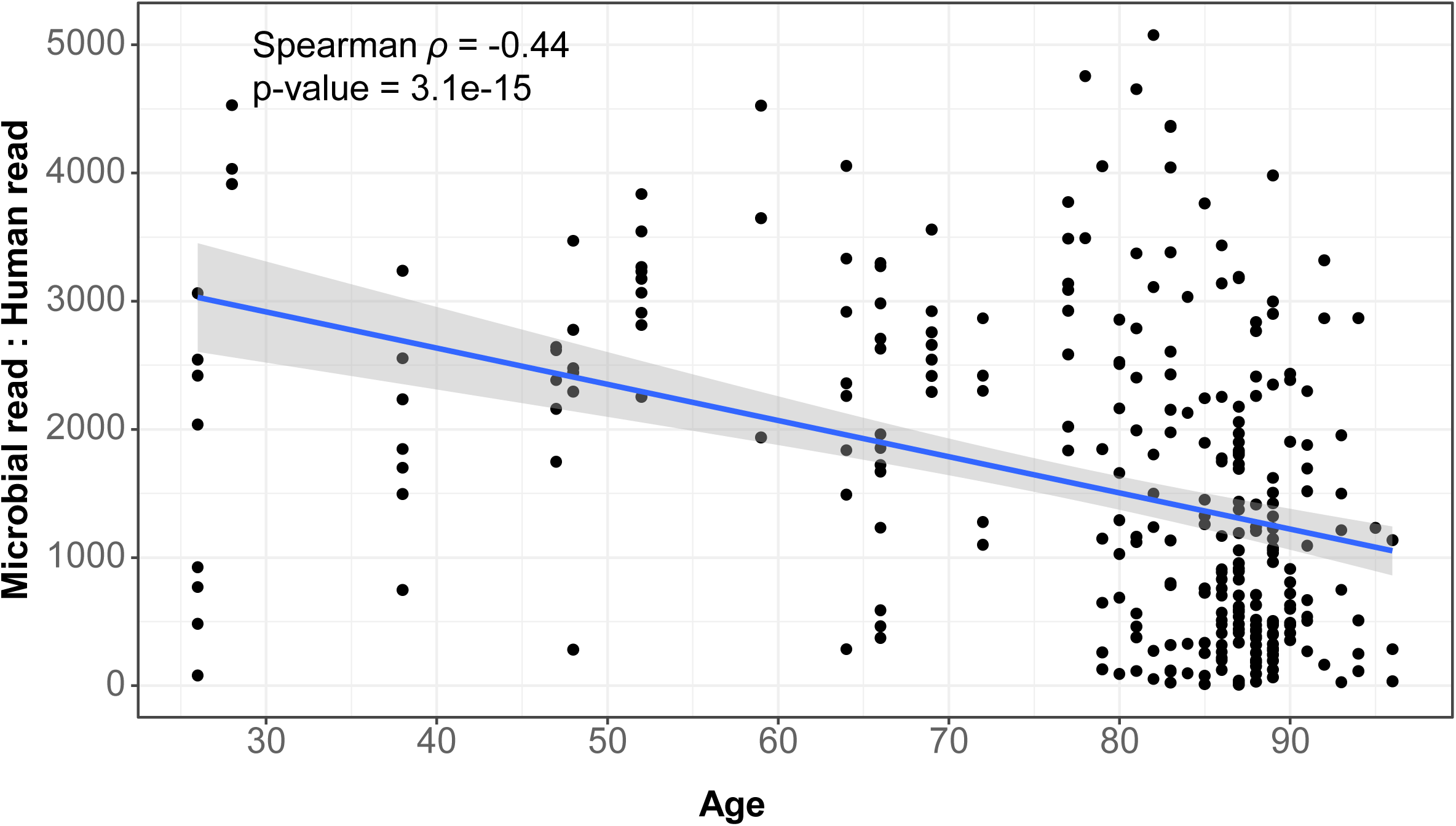
Decrease in microbe-to-human read ratio with age. Scatterplot showing microbe-to-human read ratio (as a proxy for microbial biomass compared to human biomass) in relation to the age of subjects for the SG90 and CPE cohorts. Microbe-to-human read ratio was estimated as the proportion of microbial reads relative to human reads in the dataset (minimap2 mapping). Outlier points are not included in the visualization. Regression line (in blue) shows a negative correlation of microbe-to-human read ratio with age.

**Supplementary Figure 14:**
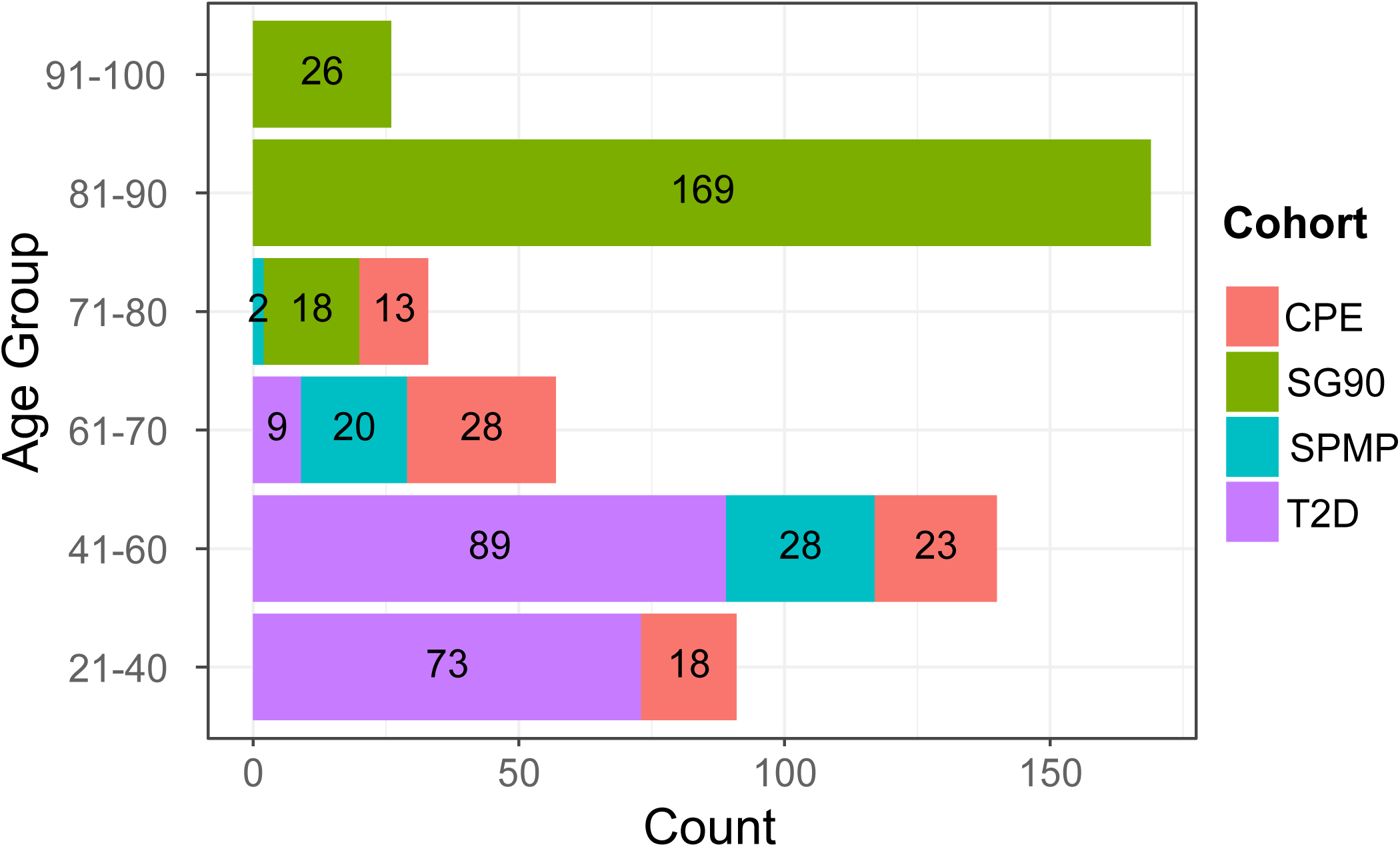
Age distribution for the different cohorts. Stacked bar chart showing the number of subjects (indicated in every bar) from different cohorts in each age groups. The number of subjects in each age group is calculated by including the lower and upper end of the range.

**Supplementary Figure 15:**
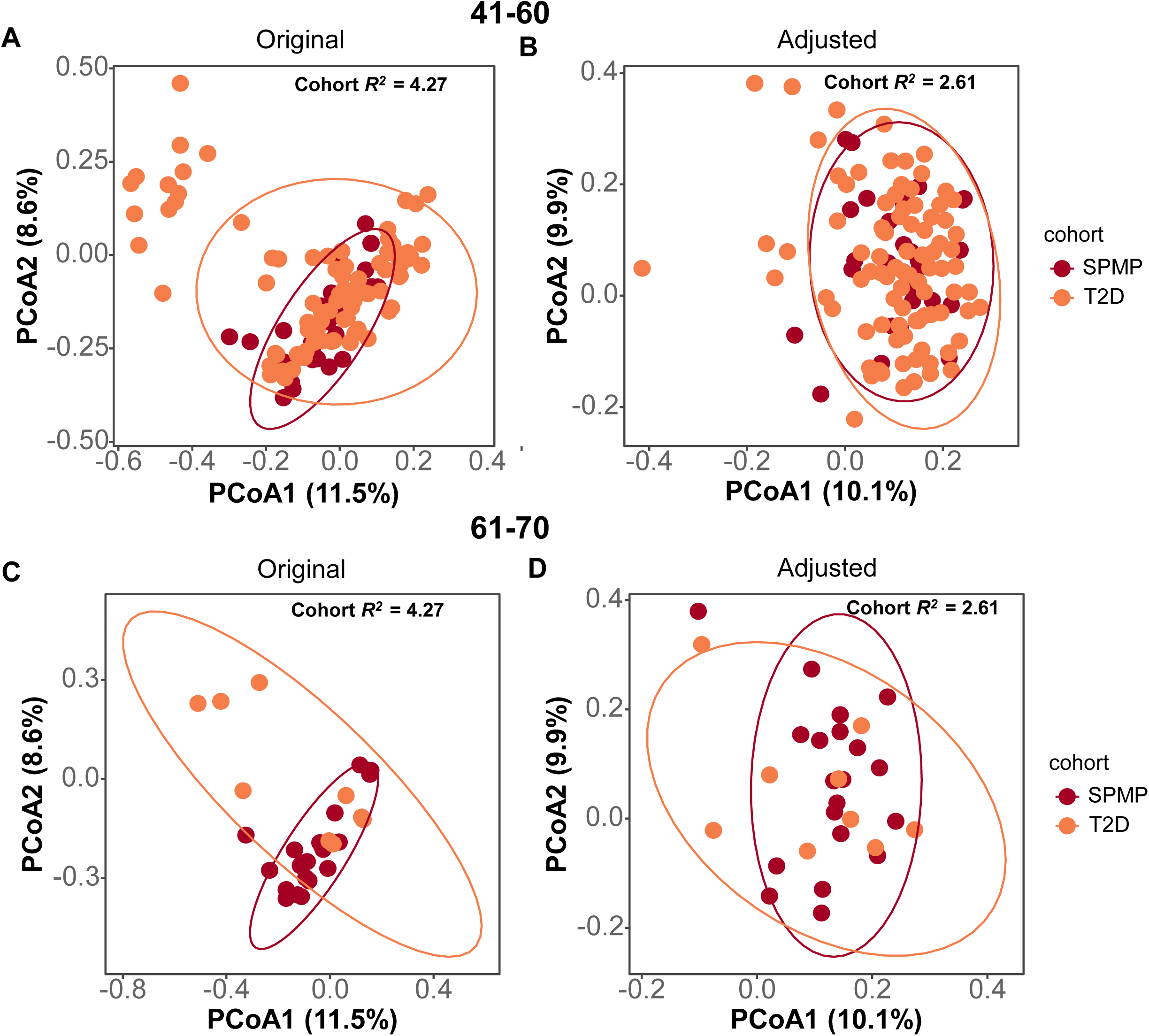
Effect of batch correction. PCoA plots using Bray Curtis dissimilarity for age-groups 41-60 (top panel) and 61-70 (bottom panel) before (A, C) and after (B, D) batch correction. For both age groups, batch correction appears to reduce the variation across cohorts as expected.

**Supplementary Figure 16:**
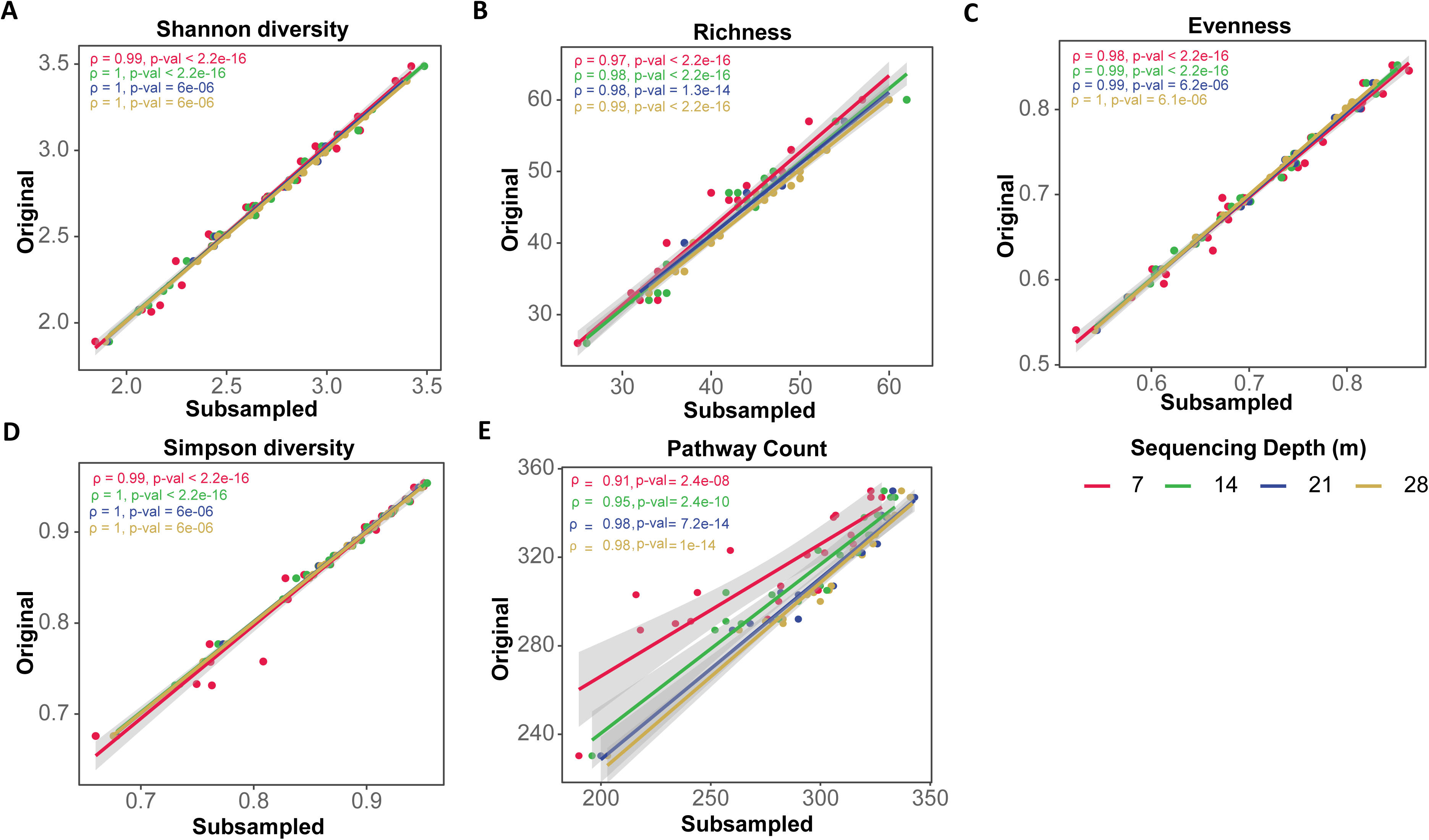
Correlation plots at different sequencing depths. Correlation plots between metrics for original datasets and subsampled sets at different sequencing depths for (A) Shannon diversity index, (B) Richness, (C) Evenness, (D) Simpson diversity index and (E) Pathway counts. Each colored line in each box represents regression line for different sequencing depths. Inset values indicate the Spearman ρ and p-values for the corresponding sequencing depths. The high Spearman correlation values show that all metrics are highly robust to variations in sequencing depth.

